# A novel interaction between MNT and CCDC6: structural insights and distinct roles in cell migration and the DNA damage response

**DOI:** 10.64898/2026.11.10.760688

**Authors:** Vanessa Junco, Elena Cabezón, Sandra Lastra, Judit Liaño-Pons, Alba Puente, M. Carmen Lafita-Navarro, Angela Celetti, Marta Martínez-Conde, Matilde Murga, Oscar Fernández-Capetillo, M. Dolores Delgado, Javier León, Lucía García-Gutiérrez

## Abstract

MNT is a transcription factor of the MYC-MAX-MXD network whose biological functions remain incompletely understood. Here, we identify CCDC6, a tumor suppressor involved in the DNA damage response (DDR), as a novel MNT-interacting protein. Co-immunoprecipitation and interaction mapping analyses identified a part of the CCDC6 coiled-coil domain and the MNT bHLH-LZ region as major determinants of binding. Structural modelling further identified MNT residues predicted to mediate CCDC6 binding that are also critical for MNT–MAX dimerization. Transcriptomic analyses revealed substantial overlap between the gene expression programs regulated by MNT and CCDC6. Loss of either protein promoted cell migration, whereas MNT and CCDC6 cooperated in the regulation of γH2AX signalling, cisplatin responses and CCDC6 localization during the DDR. Together, our findings uncover a previously unrecognized connection between CCDC6 and the MYC-MAX-MXD network through its interaction with MNT, revealing an additional layer of regulation within this transcriptional network.

## Introduction

MNT is a basic helix-loop-helix leucine zipper (bHLH-LZ) transcription factor that belongs to the MXD family, which also includes MXD1, MXI1, MXD3 and MXD4. Among these, MNT is the largest and most ubiquitously expressed. Similar to other MXD proteins, MNT forms heterodimers with MAX through their respective HLH-LZ domains to bind DNA and regulate transcription ^1,2^. MXD proteins belong to a broader network of transcription factors that includes MYC proteins. MYC is an oncogenic bHLH-LZ transcription factor frequently altered in human cancers that also forms dimers with MAX ^3,4^. MXD proteins, including MNT, compete with MYC for MAX binding. Both MNT–MAX and MYC–MAX heterodimers bind to E-boxes in the DNA, repressing or activating transcription, respectively ^5,6^. Many MYC-MAX target genes are also repressed by MNT–MAX, so MNT is often described as a functional antagonist of MYC. *Mnt*^-/-^ mice die soon after birth, while *Mxd1*^-/-^, *Mxi1*^-/-^ and *Mxd3*^-/-^ mice are viable, which suggests that *Mnt* may be the only *Mxd* gene with non-redundant essential functions ^7,8^.

MNT is the MXD protein most strongly associated with cancer. However, its role remains unclear, as it exhibits both oncogenic and tumor suppressive activities depending on the model. Tumor-suppressive functions are supported by findings showing that MNT is lost in approximately 10% of tumors ^4^, particularly Sezary syndrome, a type of cutaneous lymphoma ^9^ and that tissue-specific loss of MNT in mouse models produces thymic lymphoma ^10^, acute myeloid leukemia ^11^ or mammary tumors ^12,13^. Additionally, MNT knockdown enhances fibroblast proliferation ^12^. On the other hand, T-cell-specific homozygous *Mnt* deletion prevents thymic lymphoma in MYC-overexpressing mice ^14^, *Mnt* heterozygosity slows MYC-driven tumorigenesis ^15^ and lymphopoiesis in both wild-type and *Eμ-Myc* mice is MNT dependent ^16^. Therefore, MNT is a key component of the MYC network and has been proposed as a potential therapeutic target in cancer, although its precise functions remain to be clarified.

Only a few proteins have been reported to interact with MNT. Apart from MAX, MNT is known to interact with MAX-like protein X (MLX), another HLH-LZ protein that connects the MYC-MAX-MXD network with metabolic regulation through carbohydrate response elements ^17^; SIN3, a corepressor essential for MNT-mediated transcriptional repression ^18,19^; and REL, an NF-κB member involved in apoptosis and immune signalling pathways ^20^. Moreover, MNT is the only protein within the MYC-MAX-MXD network, apart from MAX, reported to form homodimers ^21^. Identifying new MNT protein partners is crucial to clarify its roles in physiological and pathological contexts.

We identified CCDC6 (coiled-coil domain-containing protein 6), also called PTC or H4, as a novel MNT-interacting protein. CCDC6 is a ubiquitously expressed phosphoprotein which may act as a tumor suppressor involved in the DNA damage response since it is required for proper ATM-dependent function upon genotoxic stress ^22–25^. Additionally, CCDC6 functions as a pro-apoptotic protein when phosphorylated by ERK2 ^26^. CCDC6 was originally identified in a chimeric gene between *CCDC6* and the tyrosine kinase gene *RET* generated by chromosomal rearrangement in papillary thyroid cancer, generating an oncogenic fusion protein ^27,28^. CCDC6 is also involved in rare translocations in other cancers ^29–31^. Therefore, CCDC6 has been proposed as a potential target for novel combination therapies involving tyrosine kinase inhibitors ^23,32^.

In this work, we identified and characterized a novel interaction between MNT and CCDC6 and explored its functional consequences in different cellular models. We found that a region within the CCDC6 coiled-coil domain interacts with the MNT bHLH-LZ region, suggesting that CCDC6 may represent a previously unrecognized regulator of MNT function. Biologically, MNT impaired proliferation in the presence of high levels of CCDC6. In agreement with transcriptomic analyses, loss of either protein promoted cell migration, indicating that MNT and CCDC6 converge on the regulation of migration-associated programs. Our results also revealed a closer functional interplay between both proteins during the DNA damage response. We propose that the MNT–CCDC6 interaction adds a new layer of complexity to the MYC-MAX-MXD network and contributes to the regulation of cell proliferation and the DNA damage respons

## RESULTS

### 1. MNT and CCDC6 interact through their leucine zipper domains

To identify MAX-independent MNT interactors, endogenous MNT was subjected to affinity purification–mass spectrometry (AP-MS) in MAX-deficient and MAX-expressing rat pheochromocytoma cells ^20^. This analysis identified 16 MNT-interacting proteins in MAX-deficient UR61 cells and 52 in MAX-expressing UR61 cells. Of these, five proteins—REL, CCDC6, AMPD2, QSER1, and TPP2—were found to interact with MNT irrespective of MAX expression. In a previous study, we characterized the interaction between MNT and REL ^20^. In the present work, we focused on the MNT–CCDC6 interaction because of its potential relevance to cancer. We first examined whether MNT and CCDC6 are commonly co-expressed. Analysis of data from ProteomicsDB (https://www.proteomicsdb.org/analytics/expressionHeatmap) revealed ubiquitous expression of both proteins across a panel of 38 human tissues (Supplementary Figure 1A). Co-expression of MNT and CCDC6 was further confirmed by western blot analysis in a panel of cell lines derived from different tissue origins (Supplementary Figure 1B). To validate the interaction identified by AP-MS, we examined the association between MNT and CCDC6 by endogenous co-immunoprecipitation assays in four human cell lines of different tissue origins (HEK-293T, HeLa, MCF-7, and HAP1). MNT-knockout (MNT-KO) HAP1 cells were included as a negative control. Endogenous MNT was immunoprecipitated, and its interaction with CCDC6 was confirmed by western blot analysis in three of the four cell lines tested (Figure 1A). We next assessed whether this interaction could also be detected under ectopic expression conditions. Full-length human MNT or mouse Mnt constructs tagged with Flag and HA epitopes, respectively, together with a myc-tagged human CCDC6 full length construct (Figure 1B), were transiently expressed in HEK-293T cells. Of note, mouse Mnt shares 93% identity with human MNT ^21,33^. The interaction between exogenous MNT and CCDC6 was confirmed by immunoprecipitation of MNT using either anti-HA (Figure 1C) or anti-Flag antibodies (Figure 1D), followed by western blot analysis.

**FIGURE 1.**
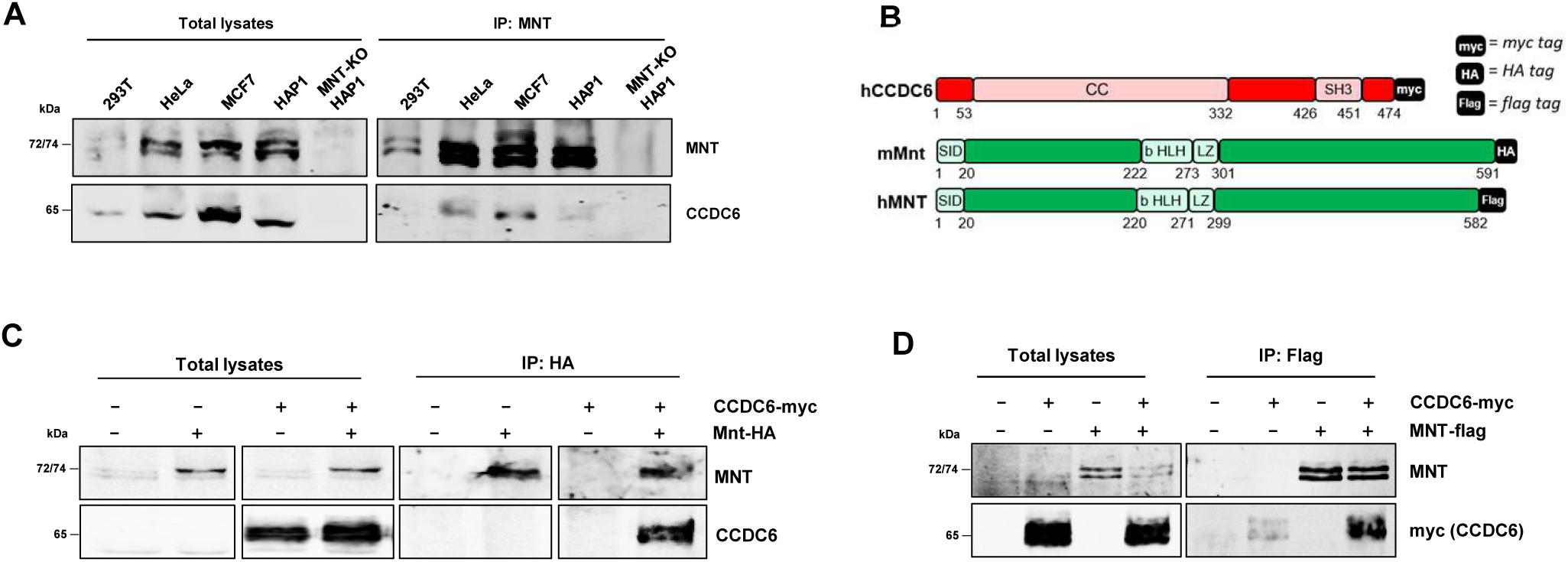
MNT interacts with CCDC6 through its N-terminal domain and LZ motif. **(A)** Co-immunoprecipitation assay of endogenous MNT and CCDC6 in different human cell lines (HEK-293T, HeLa, MCF7 and HAP1 cells). Endogenous MNT was immunoprecipitated and CCDC6 and MNT protein levels were detected by western blot. MNT-KO HAP1 cells were used as a negative control**. (B)** Schematic representation of the human CCDC6 and human and mouse MNT constructs used for co-immunoprecipitation experiments showing their respective tag sequences. **(C)** Co-immunoprecipitation assay of exogenous MNT-HA with CCDC6-myc constructs expressed in HEK-293T cells for 48 hours. Exogenous MNT was immunoprecipitated using an anti-HA antibody and levels of CCDC6 were assayed by western blot. **(D)** Co-immunoprecipitation assay of exogenous MNT-Flag with CCDC6-myc constructs expressed in HEK-293T cells for 48 hours. Exogenous MNT was immunoprecipitated using anti-Flag conjugated agarose beads and levels of CCDC6 were assayed by western blot.

To identify the regions mediating the interaction between MNT and CCDC6, we used five HA-tagged Mnt deletion mutants and two myc-tagged CCDC6 deletion mutants, schematized in Figure 2A. The Mnt mutant constructs were generated from the mouse Mnt sequence. We first determined the regions of CCDC6 required for its interaction with MNT. HEK-293T cells were co-transfected with human MNT-Flag together with full-length CCDC6-myc or the corresponding CCDC6 deletion mutants. Following immunoprecipitation of MNT using anti-Flag-conjugated agarose beads, full-length CCDC6 and the CCDC6^1–223^ mutant were found to interact with MNT, whereas the CCDC6^1–101^ mutant failed to do so (Figure 2B). These results indicate that the region spanning amino acids 101–223 of CCDC6 is required for its interaction with MNT. We next investigated the regions of MNT involved in CCDC6 binding. HEK-293T cells were co-transfected with the different Mnt-HA deletion mutants and full-length CCDC6-myc, followed by anti-HA immunoprecipitation and western blot. Analysis of N-terminal and C-terminal deletion mutants revealed that deletion of the N-terminal region abolished the interaction with CCDC6, whereas mutants lacking C-terminal sequences retained binding (Figure 2C). Specifically, the ΔNt1-Mnt mutant, which does not contain the leucine zipper (LZ) domain, failed to co-immunoprecipitate with CCDC6, while the ΔNt2– and ΔCt1-MNT and, to a lesser extent, the ΔCt2-Mnt mutants maintained the interaction, suggesting that MNT’s LZ domain and N-terminal region contribute to MNT–CCDC6 binding. To further evaluate the role of the LZ domain, HEK-293T cells were co-transfected with either wild-type or ΔLZ Mnt-HA together with full-length CCDC6-myc. Following anti-HA immunoprecipitation, CCDC6 was found to interact with both wild-type and ΔLZ Mnt with comparable efficiency (Figure 2D). While the deletion analysis points to the involvement of the LZ-containing region in MNT– CCDC6 binding, the persistence of the interaction in the absence of the LZ domain suggests that other regions of MNT also contribute to this association, including the HLH domain and/or neighbouring unstructured regions.

**FIGURE 2.**
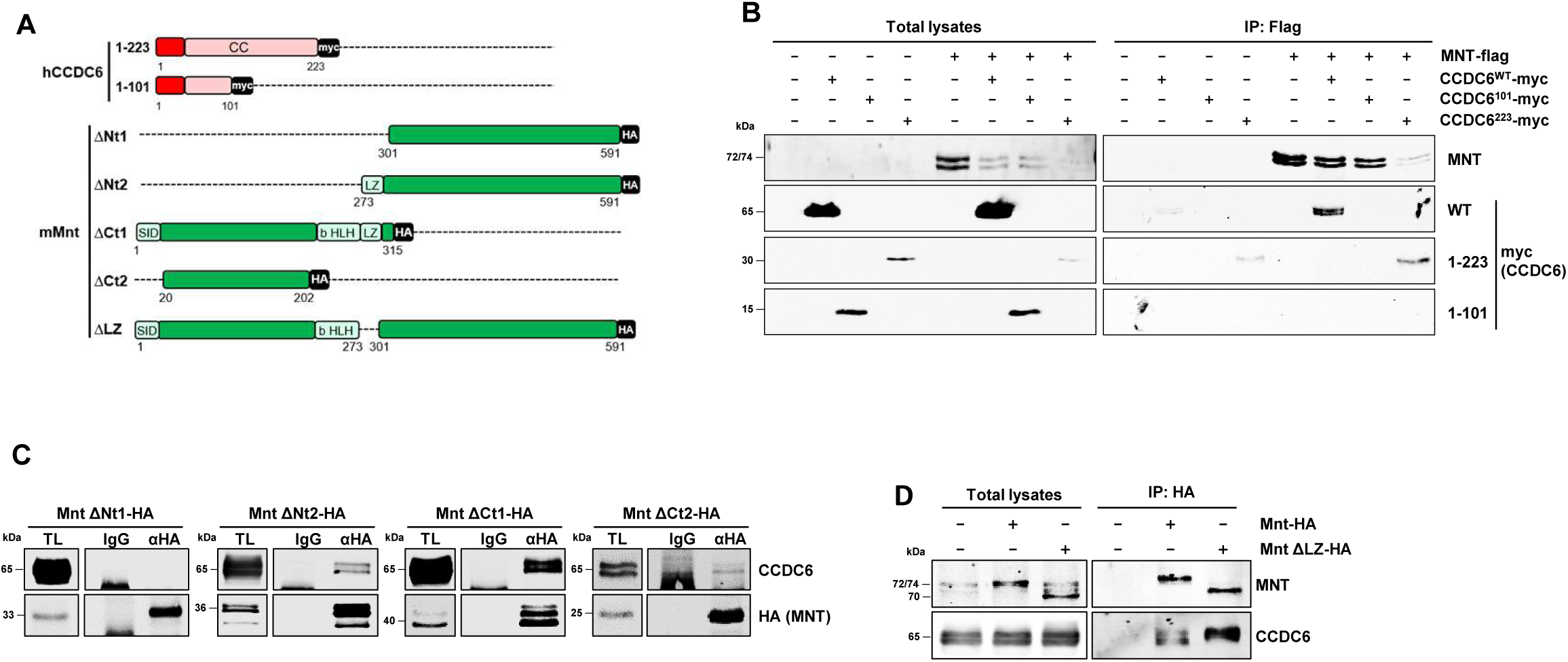
MNT interacts with CCDC6 through its N-terminal domain and LZ motif. (**A**) Schematic representation of CCDC6 and Mnt deletion mutants used in B, C and D. **(B)** Co-immunoprecipitation assays of exogenous MNT-flag with CCDC6^WT^-myc or the CCDC6^1–101^–myc or CCDC6^1–223^–myc mutants co-expressed in HEK-293T cells for 48 hours. Exogenous MNT was immunoprecipitated using anti-Flag conjugated agarose beads and levels of CCDC6 were assayed by western blot using an anti-myc tag antibody **(C)** Co-immunoprecipitation assays of CCDC6-myc with the C-terminal (ΔCt1, ΔCt2) and N-terminal (ΔNt1, ΔNt2) Mnt-HA mutants co-expressed in HEK-293T for 48 hours. Immunoprecipitation of exogenous Mnt was performed using an anti-HA antibody and CCDC6 levels were assayed by western blot. Non specific IgG was used as negative control. **(D)** Co-immunoprecipitation assays of CCDC6-myc with the wild type Mnt-HA or the ΔLZ MNT mutant co-expressed in HEK-293T for 48 hours. Immunoprecipitation of exogenous Mnt was performed using an anti-HA antibody and CCDC6 levels were assayed by western blot.

### 2. Structural modelling analyses reveal two interacting regions between MNT and CCDC6

As the regions of MNT identified by co-immunoprecipitation analyses encompassed multiple protein domains, we sought to further refine the interacting interface between MNT and CCDC6 using AlphaFold3 structural predictions ^34^. Although a large portion of MNT is predicted to be intrinsically disordered, the AlphaFold3 model of human MNT reveals a well-defined N-terminal α-helix and a structured basic helix–loop–helix leucine zipper (bHLH-LZ) domain. This domain closely resembles the corresponding region of MXD1, whose structure has been experimentally resolved by X-ray crystallography ^35^. AlphaFold3 predictions for human CCDC6 indicate two intrinsically disordered regions located at the N– and C-termini, together with two long α-helices spanning residues 53–344 and separated by a short coiled-loop region encompassing residues 235–255. Structural modelling of the MNT–CCDC6 complex predicts an interaction involving MNT residues 221–316 and CCDC6 residues 150–227 (Figure 3A, squares 1 and 2 respectively). Notably, the MNT region predicted to participate in the interaction comprises the characteristic bHLH-LZ domain, which is highly conserved among MYC, MAX, and MXD family proteins. The interacting region predicted in CCDC6 lies within its first α-helical segment and encompasses a leucine zipper motif, suggesting that this domain contributes to the interaction with MNT (Figure 3A).

**FIGURE 3.**
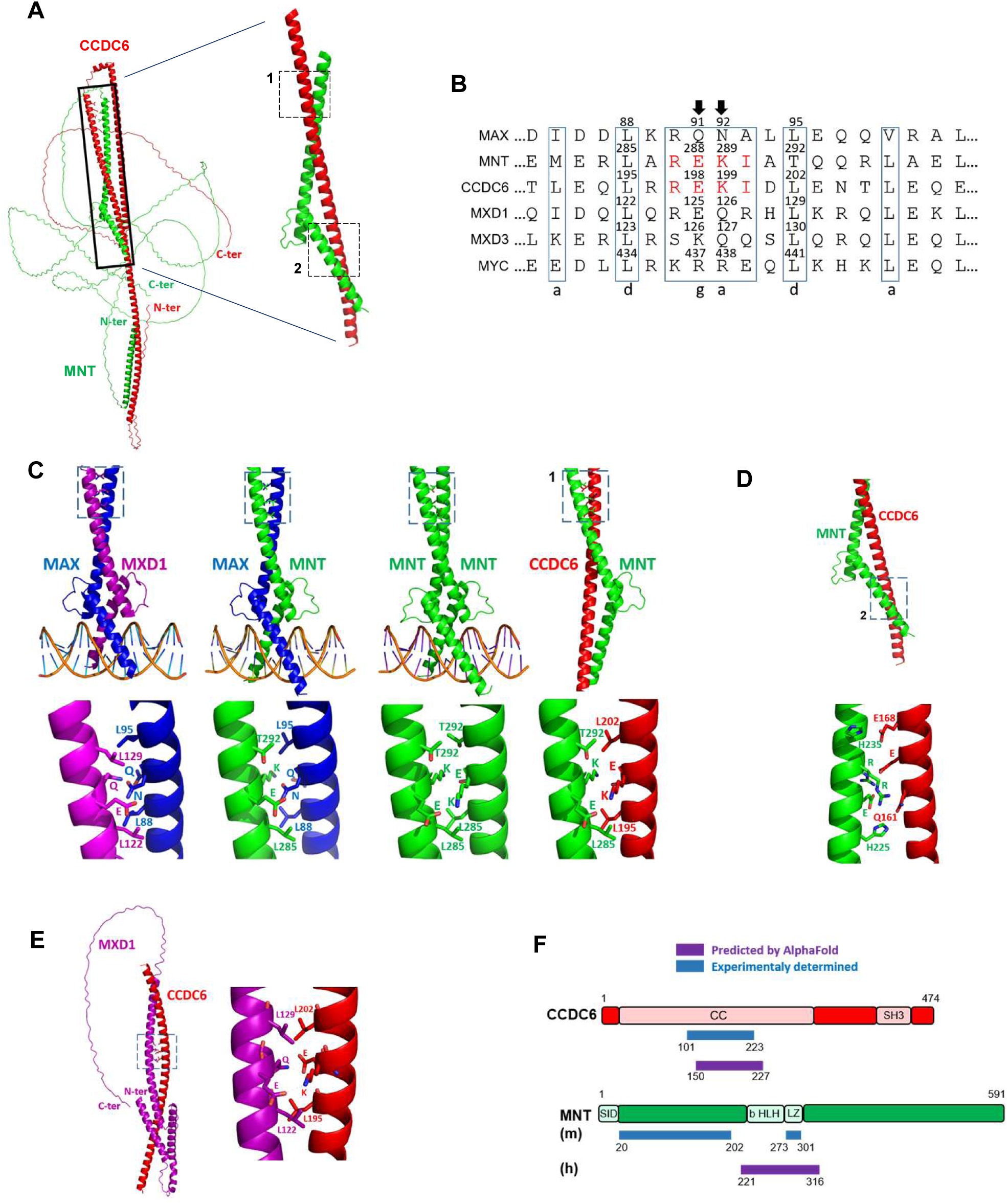
AlphaFold3 reveals a motif within the LZs of both MNT and CCDC6 that is conserved along the MYC-MAX-MXD proteins. **(A)** Prediction of MNT-CCDC6 complexes by AlphaFlold3. The dashed squares numbered 1 and 2 show the two interacting regions between MNT and CCDC6 predicted by Alphafold3. **(B)** Alignment of MAX, MNT, CCDC6, MXD1, MXD3 and MYC protein sequences (human origin) showing conserved residues involved in the formation of protein dimers. In red, the conserved regions shared by MNT and CCDC6. **(C)** Prediction of the interacting regions between MAX-MXD1, MAX-MNT, MNT-MNT and MNT-CCDC6 (interacting region 1). The conserved residues among the different members of the MYC-MAX-MXD network and CCDC6 that are involved in the different interactions are highlighted. **(D)** Prediction of the interacting region 2 between MNT and CCDC6. The important residues involved in the predicted interaction are highlighted. **(E)** Prediction of the interaction between MXD1 and CCDC6. The important residues involved in the predicted interaction are highlighted. **(F)** Comparison between the predicted MNT-CCDC6 interaction by AlphaFold3 (violet bars) and the model proposed with the results experimentally obtained by co-immunoprecipitation assays (blue bars). All analysed pdb files were downloaded from AlphaFold3 Server.

A large portion of the predicted MNT–CCDC6 interface adopts a coiled-coil conformation characterized by a heptad repeat pattern (abcdefg; Figure 3B), with hydrophobic residues occupying positions *a* and *d* within the heterodimer. Accordingly, the MNT–CCDC6 complex is predicted to be stabilized by a combination of hydrophobic and polar interactions involving two α-helices from MNT and one α-helix from CCDC6, including the leucine zipper motifs present in both proteins. Notably, several of the residues predicted to mediate the interaction with CCDC6 (interacting region 1; Figure 3A) are also involved in MNT–MAX dimerization. Thus, the first region of MNT predicted to interact with CCDC6 overlaps with the region responsible for the formation of MNT–MAX complexes. The predicted MNT–CCDC6 complex, together with the crystal structure of the MXD1–MAX heterodimer bound to an E-box ^35^, is shown in Figure 3C. We previously reported that MNT is capable of forming homodimers *in vivo* based on co-immunoprecipitation experiments ^21^, although the structural basis of this interaction was not investigated. In the resolved MXD1–MAX structure ^35^, heterodimer formation depends on interactions between MAX residues Q91 and N92 and MXD1 residues E125 and Q126 (Figure 3C). The critical glutamate residue present in MXD1 (E125) is conserved in MNT (E288). Accordingly, the residue pairs MAX Q91–N92/MNT E288– K289 are predicted to contribute to MNT–MAX heterodimerization, whereas the reciprocal interaction between MNT E288–K289 residues may underlie MNT homodimer formation. CCDC6 contains the equivalent conserved residues within the region predicted to interact with MNT (E198–K199; Figure 3B,C), suggesting that contacts between MNT E288–K289 and CCDC6 E198–K199 may constitute a key structural determinant of the MNT– CCDC6 interaction. Further analysis identified a second interaction interface between MNT and CCDC6 (interacting region 2), involving residues H225-E229-R233-H235 of MNT and Q161-E167-E168 of CCDC6 (Figure 3A,D). Interestingly, several residues within this MNT region correspond to amino acids required for DNA binding in MAX. Indeed, H28, E32 and R36 have been identified as critical for MAX–DNA interactions ^35^, and their homologous residues in MNT are H225, E229 and R233, respectively. These same residues are predicted to participate in the interaction with CCDC6, raising the possibility that binding of CCDC6 to MNT could interfere with DNA recognition (Figure 3D). Finally, we investigated whether CCDC6 could also interact with other members of the MXD family. AlphaFold3 also predicted a stable interaction between CCDC6 and MXD1 mediated by residues E125-Q126 of MXD1 and E198-K199 of CCDC6 (Figure 3E). This interaction involves E125, the conserved glutamate equivalent to MNT E288, one of the residues predicted to contribute to CCDC6 binding. In contrast, MXD3 contains a lysine residue (K126) at the position corresponding to MXD1 E125 (Figure 3B), which would be unable to establish the same interaction with CCDC6 K199. Consistent with this prediction, AlphaFold3 did not support formation of a stable MXD3–CCDC6 complex. Interestingly, AlphaFold3 did not predict a stable interaction between CCDC6 and MYC. In contrast to MNT, MYC contains two arginine residues (R437 and R438) at positions equivalent to MNT E288 and K289 (Figure 3B). The resulting electrostatic repulsion between MYC R438 and CCDC6 K199, together with a different orientation of the interacting α-helices that prevents leucine-mediated contacts, would not allow the interaction. Altogether, these observations support a preferential structural compatibility of CCDC6 with MNT and, potentially, MXD1.These structural predictions are consistent with our experimental observations, which identified the LZ-containing region of MNT and the coiled-coil region of CCDC6 as important contributors to the interaction (Figure 2C,D). Figure 3F compares the AlphaFold3-predicted MNT–CCDC6 complex with the interaction model derived from our biochemical data, revealing a strong degree of concordance between both approaches.

### 3. CCDC6 enhances cell proliferation in the absence of MNT

Having confirmed and characterized the interaction between MNT and CCDC6, we next investigated the biological functions associated with this protein complex. To this end, we used MNT– and CCDC6-knockout (KO) HAP1 and MCF7 cells generated by CRISPR-Cas9 technology (Figures 4A,B). Notably, MNT protein levels were reduced in CCDC6-KO HAP1 cells (Figure 4A), suggesting that CCDC6 may contribute, at least in part, to the stability of MNT. Given the established role of MNT in the regulation of cell proliferation, we next examined the effects of MNT or CCDC6 loss on the proliferative capacity of HAP1 and MCF7 cells. MNT depletion did not significantly alter the proliferation rate of either cell line (Figure 4C). In contrast, CCDC6 loss reduced the proliferation of HAP1 cells, whereas no appreciable effect was observed in MCF7 cells (Figure 4D). To investigate whether the effect of CCDC6 depletion on HAP1 cell proliferation was dependent on its interaction with MNT, we simultaneously modulated the expression of both proteins. CCDC6 was ectopically expressed in MNT-KO HAP1 cells by lentiviral transduction using a CCDC6-IRES-Cherry construct, while the corresponding empty vector served as a control. CCDC6 overexpression significantly increased the proliferation of MNT-KO HAP1 cells (Figure 5A,B). To further assess the functional relationship between MNT and CCDC6, MNT expression was restored in the CCDC6-overexpressing MNT-KO HAP1 cells, as well as in their corresponding control cells, by lentiviral transduction with an MNT-GFP construct. Cells transduced with the corresponding empty vector were used as controls (Figure 5C). A schematic summary of the four HAP1-derived cell populations generated for these experiments is shown in Figure 5D. Restoration of MNT expression in MNT-KO cells increased both endogenous and exogenous CCDC6 protein levels, supporting the notion that MNT contributes to CCDC6 protein stability. In turn, MNT re-expression abolished the increase in cell proliferation induced by CCDC6 overexpression in MNT-deficient cells (Figure 5E). These findings are consistent with the observation that MNT depletion alone does not affect HAP1 cell proliferation (Figure 4C), while elevated CCDC6 levels promote cell growth in the absence of MNT. Together, these results suggest that the proliferative effects of CCDC6 are influenced by MNT levels and support a model in which cell proliferation is regulated, at least in part, through the functional interaction between CCDC6 and MNT.

**FIGURE 4.**
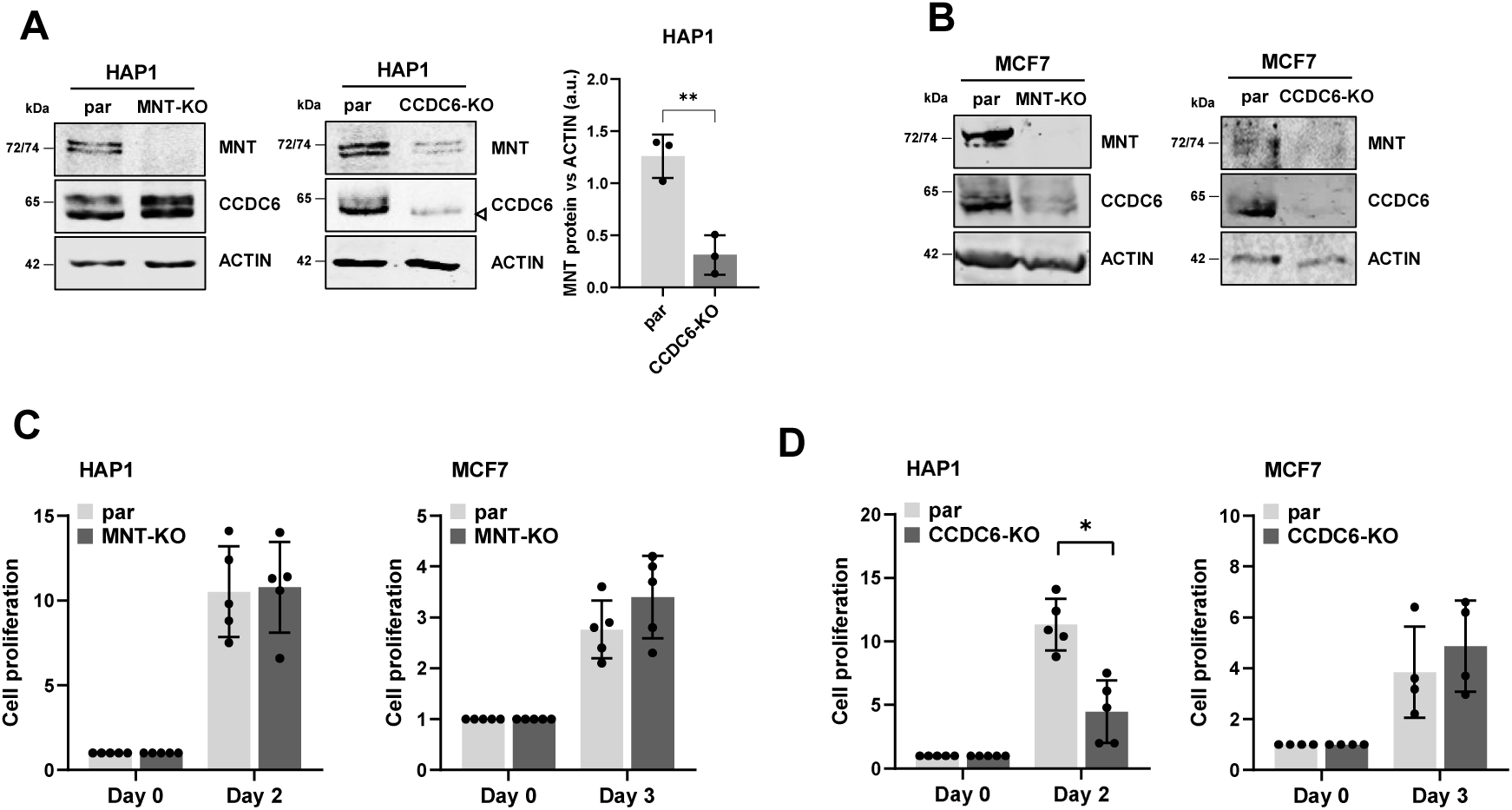
Effect of MNT or CCDC6 depletion on cell proliferation. **(A)** Validation of MNT and CCDC6 depletion in HAP1 cells by western blot. MNT and CCDC6 protein levels are shown. β-ACTIN levels were used as loading control. ◃denotes a non-specific band. The graph on the right shows the densitometric quantification of MNT protein levels relative to β-ACTIN from parental and CCDC6-KO HAP1 cells. Data are presented as mean ±SD from n = 3 independent experiments. Statistical significance was determined using a two-tailed paired t-test; ** p < 0.01. **(B)** Validation of MNT and CCDC6 depletion in MCF7 cells by western blot. MNT and CCDC6 protein levels are shown. β-ACTIN levels were used as loading control. **(C)** Proliferation assay of HAP1 or MCF7 cells comparing MNT-KO against parental cells obtained by quantification of cristal violet staining. Data are presented as mean ±SD from n = 5 independent experiments. Statistical significance was determined using a two-tailed paired t-test; ns p > 0.05. **(D)** Proliferation assay of HAP1 or MCF7 cells comparing CCDC6-KO against parental cells obtained by quantification of cristal violet staining. Data are presented as mean ±SD from n ≥ 4 independent experiments. Statistical significance was determined using a two-tailed paired t-test; * p < 0.05.

**FIGURE 5.**
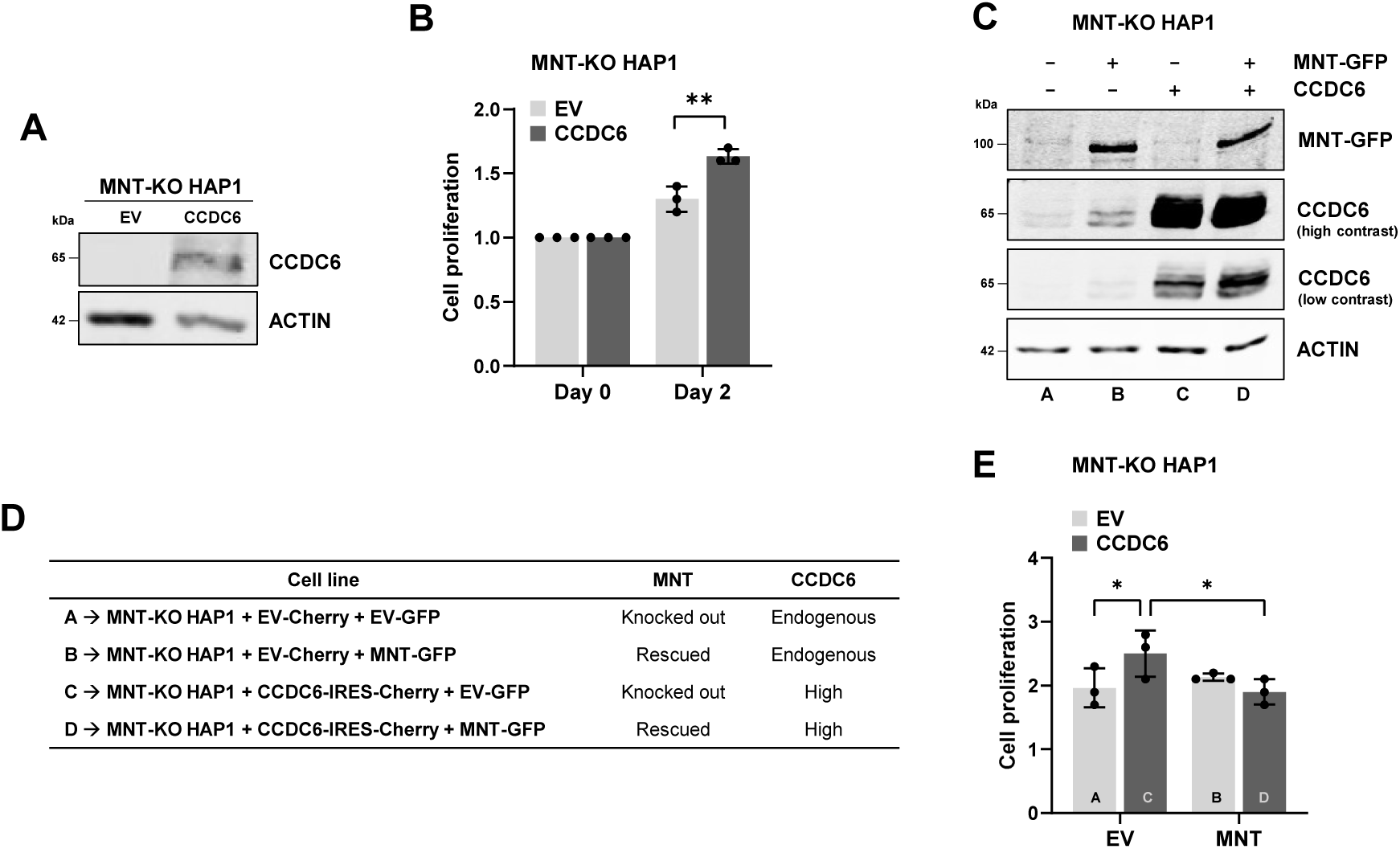
CCDC6 overexpression induces cell proliferation in the absence of MNT in HAP1 cells. **(A)** Validation of CCDC6 overexpression measured by western blot in MNT-KO HAP1 cells. CCDC6 levels are shown. β-ACTIN levels were used as loading control. **(B)** Proliferation assay of MNT-KO HAP1 cells overexpressing CCDC6 or the corresponding empty vector (EV) measured by PrestoBlue. Data are presented as mean ±SD from n = 3 independent experiments. Statistical significance was determined using a two-tailed paired t-test; ** p < 0.01. **(C)** Validation of the CCDC6 overexpression and MNT-GFP rescue in MNT-KO HAP1 cells measured by western blot. MNT-GFP and CCDC6 levels are shown. β-ACTIN levels were used as loading control. The letters at the bottom of each lane correspond to the cell lines described in D. **(D)** Summary of the MNT-KO HAP1 derived cell lines generated (i.e. CCDC6 overexpression and MNT rescue). **(E)** Proliferation assay of CCDC6 overexpressing MNT-KO HAP1 cells upon MNT-GFP rescue or its corresponding empty vector (EV) measured by PrestoBlue. The letters at the bottom of each column correspond to the cell lines described in C. Data are presented as mean ±SD from n = 3 independent experiments. Statistical significance was determined using two-way ANOVA; * p < 0.05.

### 4. Knocking out MNT or CCDC6 reveals common gene pattern alterations in HAP1 cells

To further investigate the biological processes regulated by the MNT–CCDC6 complex, we analysed bulk RNA-seq data from parental, MNT-KO, and CCDC6-KO HAP1 cells and compared the transcriptional programs associated with the loss of each protein. The RNA-seq dataset corresponding to MNT-KO HAP1 cells was previously generated by our group ^20^ while the CCDC6-KO HAP1 dataset was obtained during this work. Differentially Expressed Gene (DEG) analyses revealed that loss of either MNT or CCDC6 leads to 833 altered genes – 419 upregulated and 414 downregulated – and 1772 altered genes – 996 upregulated and 776 downregulated – respectively (log2FC ≥ 1 or ≤ –1; adj.pValue ≤ 0.05) (Supplementary Figure 2A). To validate the RNA-seq results of the CCDC6-KO HAP1, we assessed by RT-qPCR the expression of 10 representative genes (Supplementary Figure 2B). Expression changes detected in MNT-KO HAP1 cells had been previously validated ^8^. We performed Over-representation Analysis (ORA) and Gene Set Enrichment Analysis (GSEA) for both MNT-KO and CCDC6-KO HAP1 datasets (Figure 6A and Supplementary Figure 2C). The results revealed that the Epithelial to Mesenchymal Transition (EMT) Hallmark was one of the most significantly enriched pathways and that was positively regulated in both conditions (Supplementary Figure 2D). To better study whether MNT and CCDC6 share biological functions that could be explained by their protein-protein interaction, we identified the DEGs shared between MNT-KO and CCDC6-KO datasets and performed a correlation analysis. We obtained substantial overlap in the transcriptional changes induced by the loss of either protein. A total of 377 DEGs were shared between the two KO models. Among them, 149 genes were commonly upregulated and 176 genes were commonly downregulated, whereas 52 genes displayed opposite regulation in each comparison, i.e., 86.2 % of the genes were regulated in the same direction (Figure 6B, Supplementary Table 1). Correlation analysis of the 377 differentially expressed genes shared between CCDC6KO and MNTKO cells revealed a strong positive association in transcriptional changes (Pearson r = 0.71, P < 0.0001), supporting the existence of a common transcriptional programme regulated by CCDC6 and MNT. We performed ORA with the list of 377 shared DEGs and consistently with the previous analyses, the EMT Hallmark was one of the most significantly over-represented (Figure 6C). Highlighted in Figure 6B are the DEGs commonly upregulated or downregulated by MNT and CCDC6 included within the EMT Hallmark gene set (Supplementary Table 1). Among the commonly upregulated genes, *THBS1* encodes thrombospondin-1, an extracellular matrix glycoprotein involved in cell migration ^36^. Upregulation of *THBS1* was validated by RT-qPCR in both MNT-KO and CCDC6-KO HAP1 cells (Figure 6D) and was further confirmed at the protein level in HAP1 and MCF7 cells (Figure 6E,F).

**FIGURE 6.**
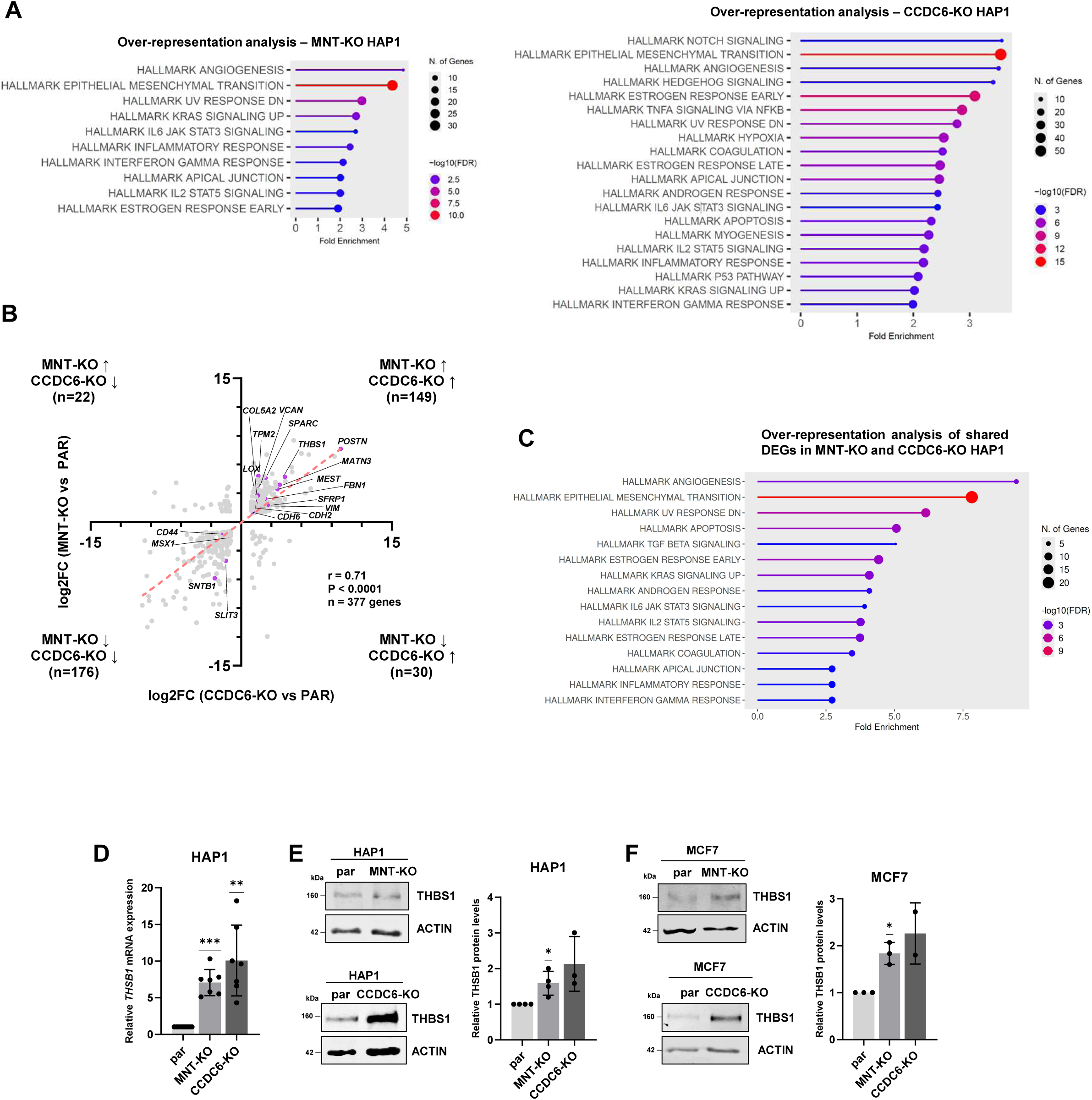
CCDC6 and MNT loss induce overlapping EMT-associated transcriptional programs. **(A)** Over-representation analysis (ORA) of Hallmark gene sets performed on differentially expressed genes (DEGs) identified in MNT-KO and CCDC6-KO HAP1 cells compared with parental (PAR) controls. The top enriched Hallmark pathways are shown according to fold enrichment, gene-set size and adjusted p-value (FDR). (**B)** Correlation analysis of DEGs shared between MNT-KO and CCDC6-KO cells. log2FC values relative to parental cells were compared for the 377 genes differentially expressed in both datasets. DEGs regulated in the same direction in both KO cell lines identified within the EMT Hallmark gene set are highlighted. The red dashed line represents the linear regression fit. Pearson correlation coefficient (r), p-value and the number of shared genes are indicated. **(C)** ORA of the 377 DEGs shared between MNT-KO and CCDC6-KO HAP1 cells. The most significantly enriched Hallmark pathways are shown according to fold enrichment, gene-set size and adjusted p-value (FDR). **(D)** *THBS1* mRNA expression measured by RT–qPCR in MNT-KO HAP1 and CCDC6-KO HAP1 cells relative to parental (Par) HAP1 cells. Data are presented as mean ±SD from n = 7 independent experiments. Statistical significance relative to the control (set to 1) was determined using a two-tailed one-sample t-test; ** p < 0.01; *** p < 0.001. **(E)** THBS1 protein levels in parental (Par), MNT-KO and CCDC6-KO HAP1 cells measured by western blot. β-ACTIN was used as a loading control. Quantification of THBS1 protein levels is shown on the right. Data are presented as mean ±SD from n ≥ 3 independent experiments. Statistical significance relative to the control (set to 1) was determined using a two-tailed one-sample t-test; * p < 0.05. **(F)** THBS1 protein levels in parental (Par), MNT-KO and CCDC6-KO MCF7 cells measured by western blot. β-ACTIN was used as a loading control. Quantification of THBS1 protein levels is shown on the right. Data are presented as mean ±SD from n ≥ 2 independent experiments. Statistical significance relative to the control (set to 1) was determined using a two-tailed one-sample t-test; * p < 0.05.

### 5. MNT and CCDC6 independently regulate cell migration

Gene set enrichment analysis identified EMT-related gene signatures among the biological processes most strongly affected by MNT or CCDC6 depletion in HAP1 cells (Supplementary Figure 2D). Given the well-established association between EMT and increased cell motility, we assessed the impact of MNT and CCDC6 loss on cell migration using *in vitro* transwell assays. Both MNT-KO and CCDC6-KO HAP1 cells exhibited increased migratory capacity compared with their parental counterparts (Figure 7A). A similar effect was observed in MCF7 cells, where depletion of either protein also enhanced cell migration (Figure 7B). As activation of the MAPK pathway is a major regulator of cell migration ^37–39^, we next examined ERK2 phosphorylation as a readout of MAPK signalling. Both MNT-KO and CCDC6-KO HAP1 cells displayed increased ERK2 phosphorylation relative to parental cells (Figure 7C,D), indicating enhanced MAPK pathway activation. Consistently, we found upregulation of *DUSP6*, a canonical ERK-responsive gene and negative-feedback regulator of MAPK signalling ^40^, together with other MAPK-associated genes among the shared DEGs obtained in the transcriptomic analyses of both knockout models, including *AXL*, *PDGFRA*, *IRS1* and *NEDD9* (Supplementary Table 1). Together, these findings reveal that loss of either MNT or CCDC6 promotes cell migration and is associated with increased MAPK signalling. To validate our findings in an independent cellular model, we took advantage of a stable shCCDC6 HeLa cell line and assessed both cell proliferation and migration. Neither CCDC6 silencing nor subsequent re-expression of CCDC6 significantly affected cell proliferation (Supplementary Figure 3A,B), consistent with the results obtained in MCF7 cells. In contrast, restoration of CCDC6 expression in shCCDC6 HeLa cells reduced their migratory capacity in *in vitro* transwell assays (Supplementary Figure 3C), further supporting a role for CCDC6 as a negative regulator of cell migration.

**FIGURE 7.**
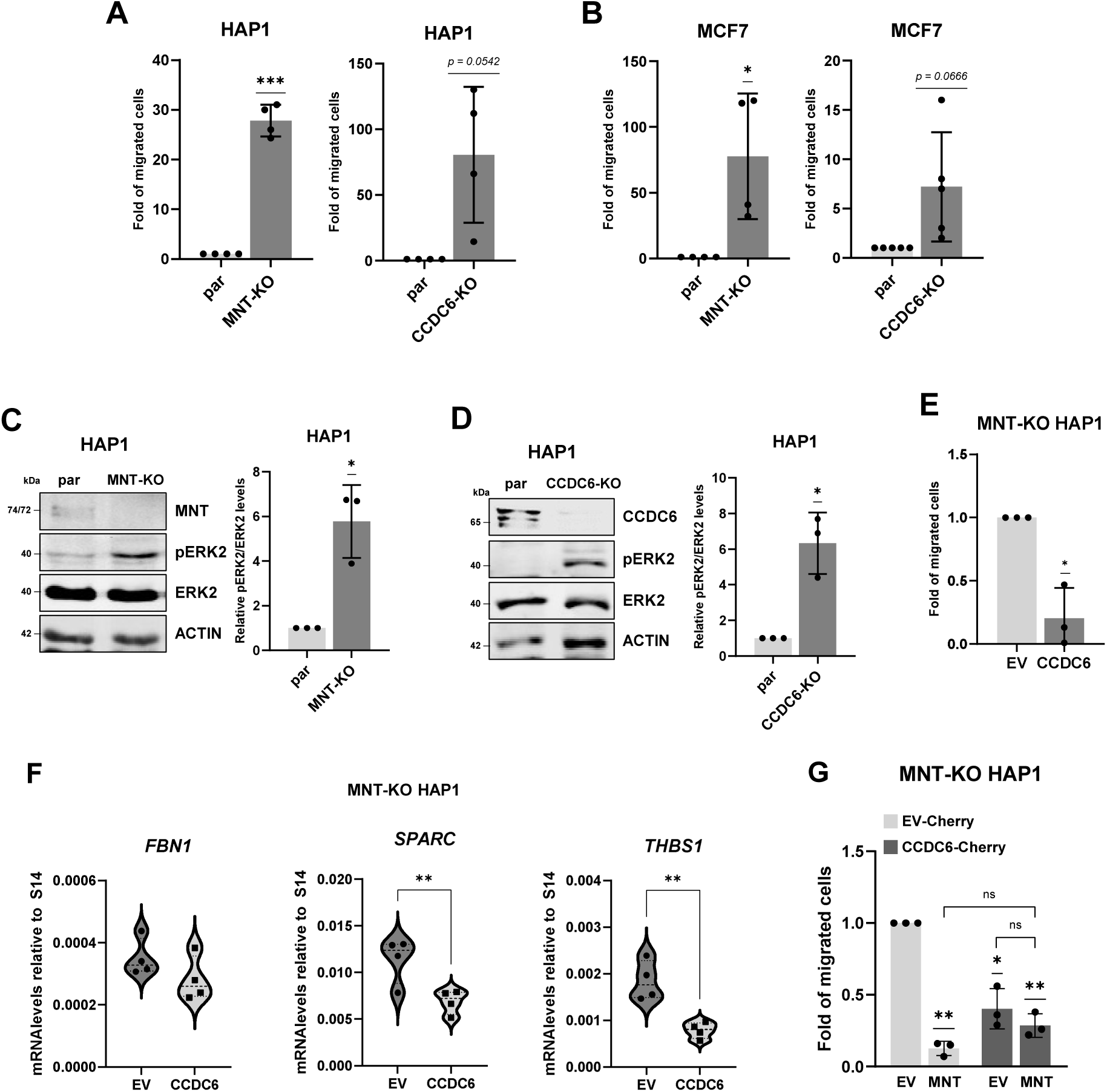
Loss of MNT or CCDC6 enhances cell migration in HAP1 and MCF7 cells. (**A**) Quantification of transwell migration assays performed with MNT-KO HAP1 and CCDC6-KO HAP1 cells relative to parental (Par) HAP1 cells. Cells were allowed to migrate for 20 h under low-serum conditions. Data are presented as mean ± SD from n = 4 independent experiments. Statistical significance relative to the control (set to 1) was determined using a two-tailed one-sample t-test; *** p<0.001. **(B)** Quantification of transwell migration assays performed with MNT-KO MCF7 and CCDC6-KO MCF7 cells relative to parental (Par) MCF7 cells. Cells were allowed to migrate for 20 h under low-serum conditions. Data are presented as mean ± SD from n ≥ 4 independent experiments. Statistical significance relative to the control (set to 1) was determined using a two-tailed one-sample t-test; * p < 0.05. **(C)** Analysis of MAPK signalling in MNT-KO HAP1 cells by western blot. MNT, phospho-ERK2 (pERK2) and total ERK2 protein levels are shown. β-ACTIN levels were used as loading control. Densitometric quantification of the pERK2/ERK2 ratio is shown on the right. Data are presented as mean ± SD from n = 3 independent experiments. Statistical significance relative to the control (set to 1) was determined using a two-tailed one-sample t-test; * p < 0.05. **(D)** Analysis of MAPK signalling in CCDC6-KO HAP1 cells by western blot. CCDC6, phospho-ERK2 (pERK2) and total ERK2 protein levels are shown. β-ACTIN levels were used as loading control. Densitometric quantification of the pERK2/ERK2 ratio is shown on the right. Data are presented as mean ± SD from n = 3 independent experiments. Statistical significance relative to the control (set to 1) was determined using a two-tailed one-sample t-test; * p < 0.05. **(E)** Quantification of transwell migration assays performed with MNT-KO HAP1 cells transiently transduced with CCDC6-Cherry or the corresponding empty vector (EV). Cells were allowed to migrate for 20 h under low-serum conditions. Data are presented as mean ± SD from n = 3 independent experiments. Statistical significance relative to the EV control (set to 1) was determined using a two-tailed one-sample t-test; * p < 0.05. **(F)** Expression levels of *FBN1*, *SPARC* and *THBS1* measured by RT-qPCR in MNT-KO HAP1 cells upon CCDC6 overexpression compared to EV. Values are normalized against *RSP14* levels. Data are presented as violin plots of n = 4 independent experiments. Statistical significance was determined using a two-tailed t-test; **p < 0.01. **(G)** Quantification of transwell migration assays performed with MNT-KO HAP1 cells stably expressing CCDC6-Cherry or the corresponding empty vector (EV). Cells were subsequently transduced with MNT-GFP or the corresponding empty vector (EV), as indicated. Cells were allowed to migrate for 20 h under low-serum conditions. Data are presented as mean ± SD from n = 3 independent experiments. Statistical significance relative to the EV control (set to 1) was determined using a two-tailed one-sample t-test; *p < 0.05, **p < 0.01. Comparisons among groups were performed using one-way ANOVA; ns, not significant (p > 0.05).

Next, we investigated whether the effects of MNT and CCDC6 on cell migration were functionally linked. To this end, we transduced MNT-KO HAP1 cells with lentiviral particles expressing CCDC6 or the corresponding empty vector and assessed their migratory capacity using *in vitro* transwell assays. CCDC6 overexpression reduced the migration of MNT-KO HAP1 cells by approximately 75% (Figure 7E), indicating that elevated CCDC6 levels can largely compensate for the absence of MNT in the regulation of cell motility. We measured the expression of three genes identified as shared upregulated DEGs in the RNA-seq analysis (*FBN1*, *SPARC* and *THBS1*) in MNT-KO HAP1 cells upon CCDC6 overexpression (Figure 7F). We found that CCDC6 overexpression resulted in the downregulation of these genes in the absence of MNT, consistent with the hypothesis that MNT and CCDC6 independently regulate migration in HAP1 cells. To further elucidate the functional relationship between both proteins, we restored MNT expression in MNT-KO HAP1 cells with or without concomitant CCDC6 overexpression using the same experimental approach described in Figure 5. As expected, re-expression of MNT significantly reduced cell migration, supporting a role for MNT as a negative regulator of this process (Figure 7G) and CCDC6 overexpression also suppressed migration in MNT-KO cells, although slightly less efficiently than MNT restoration alone (Figure 7G). Notably, re-expression of MNT in cells already overexpressing CCDC6 did not result in a further reduction in migratory capacity (Figure 7G). The observation that either MNT re-expression or CCDC6 overexpression is sufficient to restore the anti-migratory phenotype, while their combined expression does not produce an additive effect, supports the idea that both proteins operate independently within the same functional axis controlling cell motility and that this effect does not depend on MNT-CCDC6 physical interaction.

### 6. Effect of MNT on the CCDC6-mediated DNA damage response

CCDC6 plays a role in the maintenance of γH2AX signalling during DNA repair, particularly in response to DNA double-strand breaks induced by agents such as etoposide or cisplatin ^41^. Thus, we next investigated whether MNT contributes to the CCDC6-mediated DNA damage response. First, we assessed the effect of MNT or CCDC6 depletion on the sensitivity of HAP1 cells to cisplatin treatment. MNT-KO cells displayed a modest but significant increase in resistance to cisplatin, whereas CCDC6-KO cells showed a tendency toward increased sensitivity (Figure 8A). To further investigate the functional interplay between MNT and CCDC6 during the DNA damage response, we used the same MNT-KO HAP1-derived cell lines described in Figure 5D, which differ in MNT and CCDC6 expression levels. Restoration of MNT expression in MNT-KO cells increased cisplatin sensitivity (Figure 8B), indicating that MNT contributes to the cellular response to cisplatin while overexpression of CCDC6 alone had little effect. However, when CCDC6 overexpression was combined with MNT re-expression, cells became significantly more resistant to cisplatin (Figure 8B). These findings suggest that the effects of CCDC6 on the cellular response to cisplatin depend on the presence of MNT, pointing to a functional interaction between both proteins during the DNA damage response. To gain mechanistic insight into this effect, we quantified γH2AX *foci* following cisplatin treatment, given the reported role of CCDC6 in the maintenance of γH2AX levels during DNA repair. We first analysed basal γH2AX levels in untreated cells. Re-expression of MNT in MNT-KO cells did not significantly affect the number of γH2AX *foci*, whereas CCDC6 overexpression increased basal γH2AX signal, consistent with the reported role of CCDC6 in regulating γH2AX. This increase was reversed upon MNT restoration (Figure 8C). Following cisplatin treatment, CCDC6 overexpression alone did not further increase the levels of γH2AX observed in MNT-KO cells (Figure 8E and Supplementary Figure 4A). In contrast, re-expression of MNT in cells overexpressing CCDC6 resulted in a marked accumulation of γH2AX foci relative to untreated cells (Figure 8D and Supplementary Figure 4A), an effect that was not observed upon MNT rescue alone. The marked accumulation of γH2AX observed in cells expressing both MNT and high levels of CCDC6, together with their increased resistance to cisplatin (Figure 8B), suggests an increased and more effective DNA damage response in this condition. Since CCDC6 has been reported to translocate to the nucleus in response to DNA damage ^32,41^, we next examined its subcellular localization following cisplatin treatment by immunofluorescence. In MNT-KO HAP1 cells expressing exogenous CCDC6, cisplatin treatment for 24 h induced a moderate but significant accumulation of CCDC6 in the nucleus. In contrast, re-expression of MNT increased the basal nuclear localization of CCDC6, and subsequent cisplatin treatment did not significantly alter its subcellular distribution, (Supplementary Figure 4B), consistent with the increased resistance to cisplatin shown in Figure 8B. Together, these results indicate that MNT and CCDC6 jointly influence the cellular response to cisplatin and modulate γH2AX-mediated DNA damage signalling.

**FIGURE 8.**
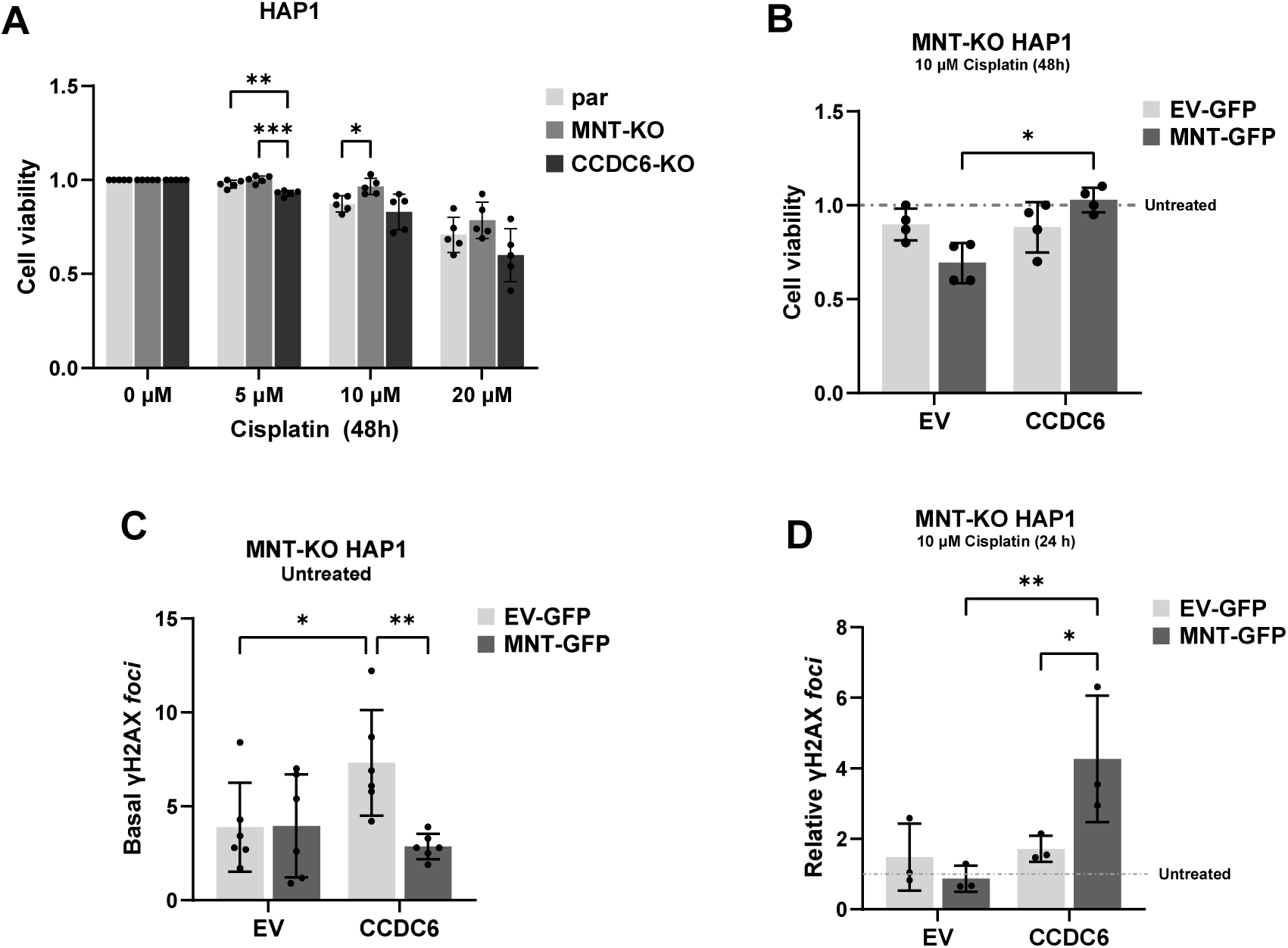
MNT and CCDC6 cooperate in the DNA damage response by modulating γH2AX signalling. **(A)** Dose-response curves of parental (par), MNT-KO and CCDC6-KO HAP1 cells treated with increasing concentrations of cisplatin. Cell viability was measured 48 h after treatment using the PrestoBlue assay. Data are presented as mean ± SD from n = 5 independent experiments. Statistical significance was determined by two-way ANOVA followed by Tukey’s multiple comparisons test; * p < 0.05, ** p < 0.01, *** p < 0.001. **(B)** Effect of MNT and CCDC6 expression on cisplatin sensitivity in MNT-KO HAP1 cells. MNT-KO HAP1 cells stably expressing CCDC6-Cherry or the corresponding empty vector (EV) were transiently transduced with MNT-GFP or the corresponding empty vector (EV). Cells were treated with 10 μM cisplatin for 48 h and viability was measured using the PrestoBlue assay. Values were normalized to the corresponding untreated controls. Data are presented as mean ± SD from n = 4 independent experiments. Statistical significance was determined by two-way ANOVA; * p < 0.05. **(C)** Basal γH2AX foci levels per nuclei in untreated MNT-KO HAP1 cells stably expressing CCDC6-Cherry or the corresponding empty vector (EV), and transiently transduced with MNT-GFP or the corresponding empty vector (EV), as indicated on the x-axis. γH2AX expression was detected by immunofluorescence and foci per nuclei quantified using ImageJ software. Five images per condition were acquired using a 40X objective. Data are presented as mean ± SD from n = 6 independent experiments. Statistical significance was determined by two-way ANOVA; * p < 0.05, ** p < 0.01. **(D)** Relative induction of γH2AX foci per nuclei in MNT-KO HAP1 cells stably expressing CCDC6-Cherry or the corresponding empty vector (EV), and transiently transduced with MNT-GFP or the corresponding empty vector (EV), as indicated on the x-axis. Cells were treated with 10 μM cisplatin for 24 h. γH2AX expression was detected by immunofluorescence and foci per nuclei quantified using ImageJ software. Five images per condition were acquired using a 40X objective. Values were normalized to the corresponding untreated controls. Data are presented as mean ± SD from n = 3 independent experiments. Statistical significance was determined by two-way ANOVA; * p < 0.05, ** p < 0.01.

## DISCUSSION

MNT is a bHLH-LZ of the MYC–MAX–MXD network with yet largely unknown functions. Unlike MYC, whose expression fluctuates throughout the cell cycle, MNT levels remain relatively constant ^33,42^, suggesting that its activity is primarily controlled through post-translational modifications and protein–protein interactions. Although MNT is one of the most extensively studied MXD family members due to its recurrent loss in several cancer types, relatively little is known about its interacting partners. Besides MAX, MNT interacts with MLX ^6^, and c-REL ^20^ and is unique among MXD family members in its ability to form homodimers ^21^. To further expand our understanding of MNT biology beyond the MYC–MAX–MXD network, we previously identified CCDC6 as a potential MAX-independent MNT interactor through proteomic analyses ^21^.In the present study, we validated the interaction between MNT and CCDC6 in different cell lines from distinct tissue origins, indicating that this interaction is conserved. Mapping analyses revealed that MNT binds a region of CCDC6 spanning amino acids 101–223. This finding is particularly relevant in cancer, as most oncogenic CCDC6 rearrangements retain only the first 101 amino acids of CCDC6 ^26^. Our results therefore predict that the majority of CCDC6 fusion proteins described in human tumours have lost the ability to interact with MNT, whereas those retaining sequences beyond residue 223 may preserve this interaction. Analysis of MNT deletion mutants indicated that both its leucine zipper (LZ) domain and, to a lesser extent, the N-terminal region of MNT contribute to CCDC6 binding. Consistent with these findings, structural modelling predicted an interaction between the coiled-coil region of CCDC6 and the two helices of the MNT bHLH-LZ domain. The predicted MNT residues overlap with those required for MNT–MAX dimerization (T292, K289, E288 and L285). The interacting CCDC6 residues (L195, E198, K199 and L202) closely resemble the MAX residues involved in interactions with MNT, MYC and other MXD proteins. The consistency between the experimental mapping data and the structural model strengthens the evidence supporting the proposed MNT– CCDC6 interaction region. Of note, the interaction detected with the MNT ΔCt2 mutant was considerably weaker than that observed with other constructs (Figure 2C) and was not supported by AlphaFold3 modelling. Thus, while additional interaction surfaces outside the bHLH-LZ domain cannot be excluded, our data suggest that the bHLH-LZ region most likely constitutes the major determinant of MNT binding to CCDC6. We also observed that CCDC6 depletion reduced MNT protein levels, whereas restoration of MNT in MNT deficient cells increased CCDC6 levels. These findings are consistent with the physical interaction between the two proteins and suggest a possible reciprocal regulation of protein stability. If confirmed, alterations in the expression of one protein in cancer could indirectly affect the abundance and function of the other. Part of the predicted MNT–CCDC6 interface overlaps with the surface used by MNT to interact with MAX, raising the possibility that MNT–CCDC6 and MNT–MAX complexes may be dynamically regulated. Despite the structural similarity between CCDC6 and MAX within this interaction region, AlphaFold3 did not predict stable interactions between CCDC6 and MYC. This lack of compatibility may be explained by the presence of two arginine residues in MYC (R437 and R438) at positions corresponding to MNT E288 and K289, generating an unfavourable electrostatic environment for interaction with CCDC6 K199. In addition, the relative orientation of the interacting α-helices differs from that observed in the MNT–CCDC6 complex, preventing the formation of stabilizing leucine-mediated contacts. These structural features resemble those proposed to explain the inability of MYC to form homodimers ^43^. Instead, preferential binding was predicted for MNT and MXD1, but not for MXD3. Structural modelling also identified a second potential interaction region involving the second α-helix of the MNT HLH domain. This region contains residues homologous to those required for DNA binding in MAX and, consistently, the structural model did not support a stable interaction between the MNT–CCDC6 complex and the DNA. This raises the possibility that CCDC6 binding could interfere with MNT’s DNA recognition. Experimentally, an MNT mutant lacking only the LZ domain retained the ability to bind CCDC6, indicating that the LZ is enough but not strictly required for the interaction. This finding is consistent with the AlphaFold3 model, which predicts an additional interaction interface involving residues within the second α-helix of the HLH domain that are preserved in the ΔLZ mutant.

Transcriptomic analyses from MNT-KO and CCDC6-KO cells revealed a substantial overlap between the transcriptional programs controlled by MNT and CCDC6 (86.2 % of DEGs), which was consistent with a role of the MNT-CCDC6 complex in regulating specific biological functions. EMT-related signatures were significantly enriched in both knockout models and, in agreement, depletion of either MNT or CCDC6 increased migration in HAP1 and MCF7 cells *in vitro*. Since MNT and CCDC6 have been proposed to act as tumour suppressors in multiple cancer types, their loss may contribute to tumour progression by promoting migratory and invasive phenotypes. Loss of either protein also increased activation of the MAPK signalling pathway, which suggests a potential contribution of this pathway to the cellular changes associated with MNT or CCDC6 loss ^37–40^. Our functional data do not support an obligate role for the MNT–CCDC6 interaction in the regulation of migration. Both proteins control a common set of EMT-and migration-associated genes, but their anti-migratory activities do not require their physical interaction, pointing towards parallel and partially independent modes of action. In fact, CCDC6 overexpression strongly reduced migration and regulated the expression of shared EMT-related genes independently of MNT. Moreover, re-expression of MNT in CCDC6-overexpressing cells did not significantly enhance this effect on migration, whereas a greater effect might be expected if it depended on the interaction between both proteins. However, the effects of MNT and CCDC6 on proliferation differ and seem to be uncoupled from their effects on migration. MNT loss did not affect proliferation in any of the cell lines tested, whereas CCDC6 depletion reduced proliferation in HAP1 cells but not in MCF7 cells. The role of CCDC6 in proliferation remains controversial, as both pro– and anti-proliferative effects have been reported ^41,44^. GSEA of CCDC6-KO cells also displayed a decrease in MYC activity, shown by the significant negative enrichment of the MYC_TARGETS_V1 Hallmark gene set and other metabolism-related pathways (Supplementary Figure 2B), which fits with their reduced proliferation with respect to their wild-type counterparts. On the other hand, CCDC6 overexpression in MNT-deficient cells increased proliferation while reducing migration, and the proliferative effect was abolished upon MNT re-expression. Thus, the proliferative effects of CCDC6 depend on MNT levels in our model. The ability of MNT to restrain CCDC6-driven proliferation is consistent with the reduced expression of MYC target genes observed in CCDC6-KO cells, suggesting a potential functional connection between CCDC6 and the MYC–MAX–MXD transcriptional network. However, our data do not address this possibility and it would require further validation.

Given the established role of CCDC6 in the DNA damage response through the stabilization of γH2AX and the maintenance of DNA damage signalling ^22–25^, we investigated the contribution of MNT to this process. MNT is a major antagonist of MYC, a well-known driver of replication stress and genome instability ^45,46^, suggesting that deregulated MYC-dependent programs may contribute to chronic activation of DNA damage signalling in the absence of MNT. Analysis of γH2AX further supports this notion. CCDC6 overexpression increased basal γH2AX levels in MNT-deficient cells, which likely reflects the stabilization of pre-existing γH2AX foci, whereas MNT re-expression reverted this effect. These observations support the idea that MNT may protect from endogenous DNA damage, in part, through proper CCDC6-mediated damage-induced signalling. The effects observed following cisplatin treatment differed from those detected under basal conditions. CCDC6 overexpression alone did not further increase γH2AX accumulation or alter cell survival in MNT-deficient cells. In contrast, the highest levels of γH2AX were observed in cells expressing both MNT and elevated CCDC6, which also showed increased resistance to cisplatin. The marked accumulation of γH2AX following cisplatin treatment likely reflects more efficient DNA damage signalling and repair. CCDC6 depletion slightly increased the sensitivity of HAP1 cells to cisplatin, in agreement with previous studies ^41^. In contrast, loss of MNT conferred moderate but significant resistance, which was reversed upon MNT re-expression. To our knowledge, the involvement of MNT in the DNA damage response has not been previously reported. Interestingly, this resistance phenotype was abolished in the presence of high CCDC6 levels, suggesting that the cellular response to cisplatin depends on the relative abundance of both proteins. Besides, CCDC6 underwent significant nuclear translocation following cisplatin treatment in MNT-deficient cells, whereas in the presence of MNT it was already predominantly nuclear and its localization was less affected by the treatment. This observation suggests that MNT may facilitate the nuclear availability of CCDC6 under basal conditions, thereby priming cells for a more efficient DNA damage response. The physical interaction between MNT and CCDC6 is consistent with the functional interplay observed during the DNA damage response, although whether direct MNT–CCDC6 binding is required for the regulation of γH2AX dynamics remains to be determined. Together, our results identify the MNT–CCDC6 axis as an important regulator of DNA damage signalling and cellular adaptation to genotoxic stress.

Taken together, our results identify a previously unrecognized connection between CCDC6 and the MYC–MAX– MXD network through its interaction with MNT. Given the partial overlap between the MNT regions involved in CCDC6 and MAX binding, it will be important to determine whether CCDC6 modulates the assembly, DNA-binding capacity or transcriptional activity of MNT-containing complexes. Such a mechanism could link DNA damage signalling to the regulation of MNT-dependent gene expression, among other effects, thereby expanding the functional repertoire of both CCDC6 and the MYC–MAX–MXD network.

## MATERIALS AND METHODS

### Cell lines and cell proliferation assays

The cell lines used in this study and their corresponding culture media are listed in Supplementary Table 2. Cell proliferation was assessed either by crystal violet staining or using the PrestoBlue Cell Viability Reagent (Invitrogen) according to the manufacturer’s instructions. For crystal violet assays, cells were stained with 0.5% crystal violet in 20% methanol for 15 min, washed extensively with water, and air-dried. For quantification, the bound dye was solubilized in 10% acetic acid and absorbance was measured at 590 nm using a GloMax® microplate reader (Promega). For PrestoBlue assays, 5 × 10^3^ cells per well were seeded in 96-well plates. At the indicated time points, PrestoBlue reagent was added and cells were incubated at 37°C. Fluorescence was measured using a GloMax® microplate reader (Promega) equipped with the Green Optical Kit (Promega). Where indicated, cells were treated with cisplatin (Sigma-Aldrich) at the specified concentrations and times. DMSO was used as vehicle control.

### Cell migration assays

Cell migration was assessed using transwell chambers with 8-μm pore-size inserts (Corning). A total of 3 × 10^4^ cells suspended in 200 μL of serum-free medium were seeded into the upper chamber, while 600 μL of complete medium was added to the lower chamber as a chemoattractant. Cells were allowed to migrate for 20 h at 37°C. Following incubation, inserts were washed with PBS and stained with crystal violet. Non-migrated cells remaining on the upper surface of the membrane were carefully removed with a cotton swab, whereas cells that had migrated to the lower surface were imaged using a Nikon Eclipse TS100 microscope. Five random fields per insert were photographed at 10× magnification, and migrated cells were quantified using ImageJ software.

### Lentiviral production and transduction

Lentiviral particles were produced in HEK293T cells by PEI-mediated co-transfection of the transfer vector with psPAX2 and pCMV-VSV-G plasmids. Viral supernatants were collected at 48 and 72 h, filtered, concentrated by PEG8000 precipitation, and stored at −80°C. Viral titers were determined in HeLa cells. HAP1, HeLa, and MCF7 cells were transduced at a MOI of 3-5 in the presence of Polybrene and, when required, subjected to antibiotic selection 48 h post-infection. Plasmids and lentiviral constructs are listed in **Supplementary Table 3**.

### Generation of MNT-KO and CCDC6-KO MCF7 cell lines

Guide RNA (gRNA) sequences used to generate human MNT and CCDC6 knockout MCF7 cell lines were selected from previously validated targets available from Horizon Discovery (HAP1 knockout cell lines). The target sequences were: CCDC6, 5′-CGCTGGCTTTGCGCAGGTCG-3′, and MNT, 5′-TCCCCTACTGCCGGACTCGA-3′. To facilitate cloning into the CRISPR/Cas9 vector, complementary oligonucleotides containing the appropriate overhangs were designed and annealed. The resulting double-stranded inserts were cloned into the BsmBI site of the lentiCRISPRv2GFP plasmid (Addgene plasmid #82416). Lentiviral particles expressing the corresponding gRNAs were produced in HEK293T cells as described above. MCF7 cells were subsequently transduced with the viral supernatants, and successfully transduced cells were enriched by fluorescence-activated cell sorting (FACS) based on GFP expression.

### RNA purification and RT-qPCR

RNA extraction, cDNA conversion and RT-qPCR analysis was performed as previously described ^20^. Briefly, total RNA was extracted using NZYol reagent (NZYTech) following the manufacturer’s instructions. cDNA was synthesized with the iScript cDNA Synthesis Kit (Bio-Rad), and gene expression was quantified by RT-qPCR using PowerUp SYBR Green Master Mix (Thermo Fisher Scientific) in a CFX Connect Real-Time PCR System (Bio-Rad). Transcript levels were normalized to *RPS14* expression. Primer sequences are listed in Supplementary Table 4.

### RNA-seq analysis

Total RNA was extracted using the RNeasy Kit (Qiagen) RNA extraction kit. Three biological replicates were analysed for each experimental condition. Gene expression counts were generated using the DRAGEN RNA pipeline (Illumina). Differential expression analysis and Gene Set Enrichment Analysis (GSEA) were performed in iDEP, and over-representation analysis (ORA) was conducted using ShinyGO v0.86.0, both available at https://bioinformatics.sdstate.edu/. Hallmark.MSigDB was used as the pathway database for enrichment analyses. Genes with a log₂ fold change ≥ 1or ≤ –1 and p-value < 0.05 were considered differentially expressed.

### Western blot and immunoprecipitation assays

Total protein extracts and immunoprecipitation assays followed by western blot analysis were performed as previously described ^20^. For total protein extraction, cells were lysed in RIPA buffer, whereas a milder lysis buffer (50 mM Tris-HCl, pH 7.5; 150 mM NaCl; 1% NP-40; 1 mM EDTA, pH 8.0; and 0.5 mM EGTA, pH 8.0) was used for immunoprecipitation experiments. Protease and phosphatase inhibitors were included in all lysis buffers. For immunoprecipitation of endogenous proteins, cell lysates were incubated with the corresponding antibodies overnight at 4°C. For immunoprecipitation of ectopically expressed tagged proteins, antibody incubation was performed for 2 h at 4°C. Immune complexes were captured using Protein G-conjugated Dynabeads™ magnetic beads (Invitrogen), washed three to four times with immunoprecipitation lysis buffer, and eluted prior to SDS-PAGE analysis. For the immunoprecipitation of flag-tagged proteins, M2 anti-Flag-conjugated agarose beads (Sigma-Aldrich) were used. Membranes were probed with IRDye® 680– or IRDye® 800-conjugated secondary antibodies (LI-COR Biosciences), and fluorescent signals were detected using an Odyssey infrared imaging system (LI-COR Biosciences). Densitometric analysis of western blot signals was performed using ImageJ software. Protein expression levels were normalized to the corresponding loading control and are presented as relative values. Antibodies used are listed in Supplementary tables 5 and 6.

### Immunofluorescence assays

Protein detection by immunofluorescence was performed as previously described ^20^. Briefly, cells were grown on glass coverslips and, following the indicated treatments, were fixed with 4% paraformaldehyde. Samples were subsequently permeabilized, blocked, and incubated with the appropriate primary antibodies overnight at 4°C. After washing, cells were incubated with FITC-conjugated secondary antibodies (Jackson ImmunoResearch Laboratories). Nuclei were counterstained with Hoechst 33342.

Fluorescence images were acquired using a Zeiss Axio Imager M1 microscope. Image processing and quantitative analyses were performed using ImageJ software (https://imagej.nih.gov/ij/). Antibodies used are listed in Supplementary tables 5 and 6.

### Structural Analysis of AlphaFold3-predicted models

All analyzed pdb files were downloaded from AlphaFold3 Server ^34^. PyMOL (https://www.pymol.org) was used to visualize the structural models and prepare the figures. Protein-protein interactions were analysed using PDBePISA (https://www.ebi.ac.uk/pdbe/pisa/pistart.html) ^47^. For structural comparison, the crystal structure of the MXD1–MAX complex bound to DNA (PDB 1NLW) was used.

### Statistical analyses

Statistical analyses were performed using GraphPad Prism version 10.6.1 for Windows, (GraphPad Software, Boston, Massachusetts USA). The specific analysis used to assess the statistical significance of each experiment is specified in the corresponding figure legend. Bars represent the mean ± standard deviation (SD) and “n” values correspond to the number of independent experiments included. GraphPad style (GP) was used to represent P values: * p<0.05; ** p<0.01; *** p<0.001; **** p<0.0001. P values ≥ 0.05 were considered not significant.

### Funding

This research was supported by grants PID2020-115903GB-I00, funded by MCIN/AEI/10.13039/501100011033 (Gobierno de España) and “FEDER, Una manera de hacer Europa” (European Union), to J.L. and M.D.D., and by the Contrato Programa Gobierno de Cantabria–Universidad de Cantabria (55.VQ10.64662) to M.D.D. and J.L.

## Acknowledgments

We are grateful to Ignacio Varela for critical reading of the manuscript and valuable advice, and to Mónica Molina for assistance with the RNA-seq data analysis. V.J.R. was supported by a PhD fellowship from the Universidad de Cantabria. L.G.G. was the recipient of a María Zambrano contract (RMZ-14) from the Universidad de Cantabria and the Ministerio de Universidades, funded by the European Union (NextGenerationEU) through the Plan de Recuperación, Transformación y Resiliencia.

## Data availability

All data supporting the findings of this study are available within the article and its Supplementary Information. The RNA-sequencing datasets generated during this study will be deposited in a public repository and released upon publication. Before publication, access to the datasets can be obtained from the corresponding author upon reasonable request.

---

**SUPPLEMENTARY FIGURE 1:**
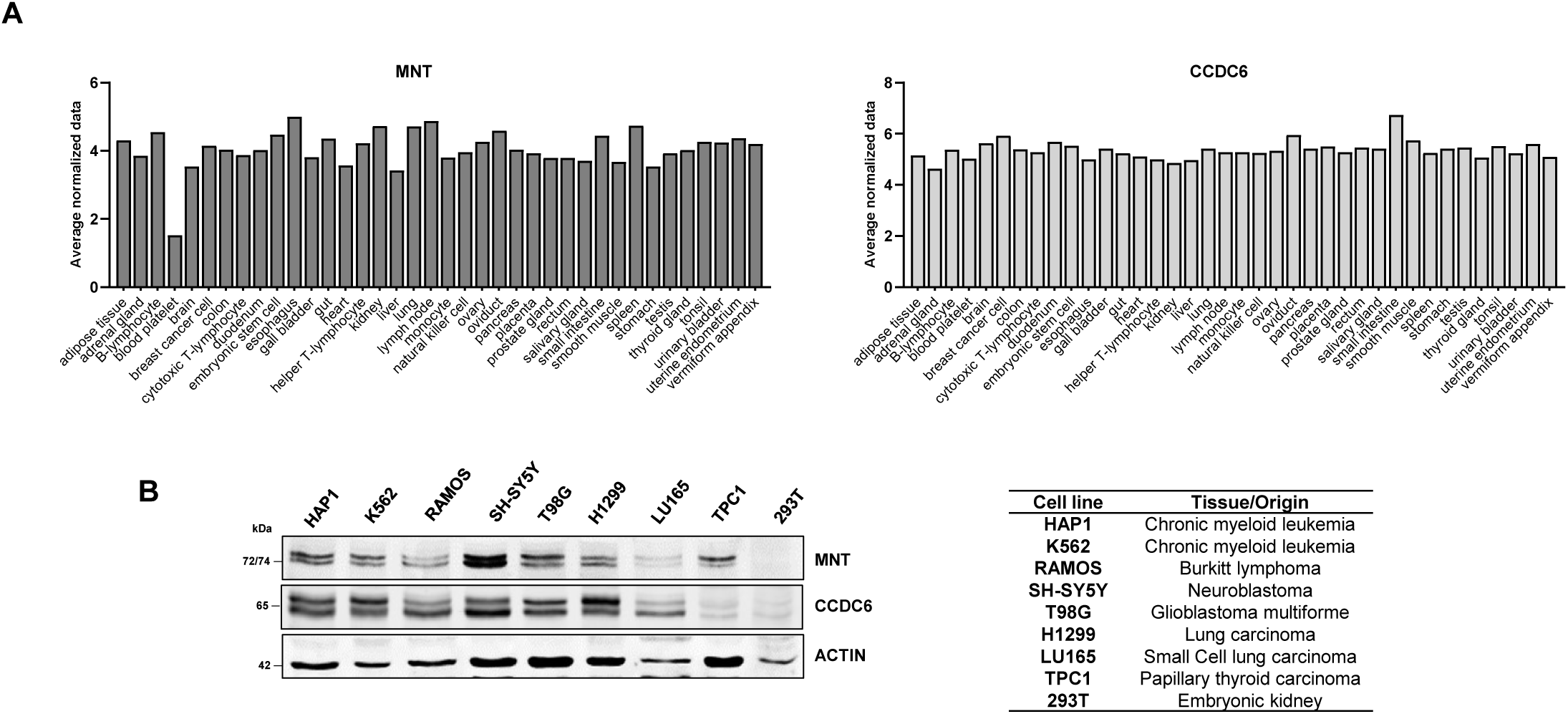
Protein expression of MNT and CCDC6 across tissues and cell lines. (**A**) Protein expression levels of MNT (letf) and CCDC6 (right) across different tissues determined by mass spectrometry. Data obtained from ProteomicsDB (https://www.proteomicsdb.org/protein). (**B**) MNT and CCDC6 protein levels across 9 different human cell lines of different origin determined by western blot. β-ACTIN levels were used as loading control.

**SUPPLEMENTARY FIGURE 2.**
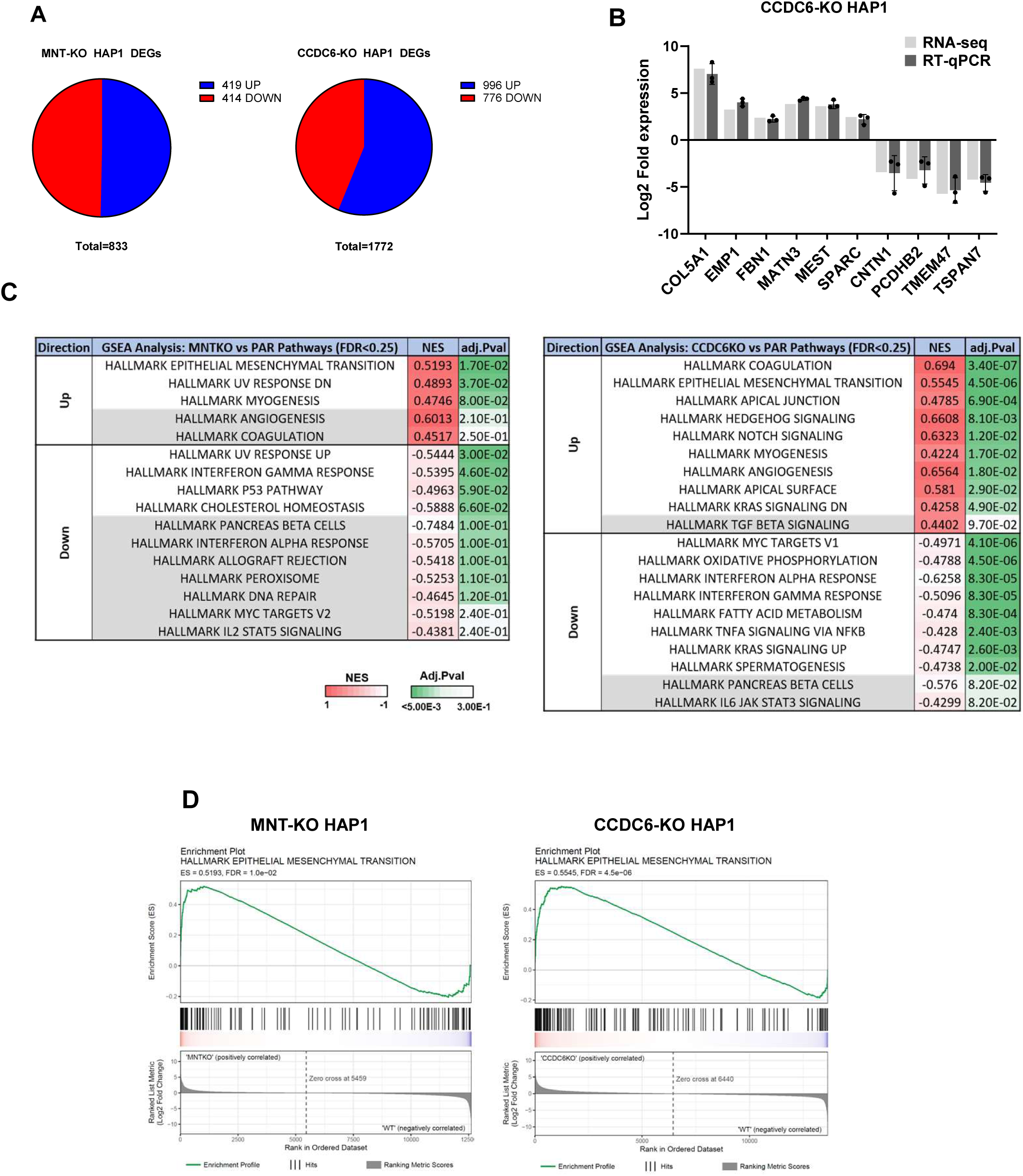
Transcriptomic profiling of MNT-KO and CCDC6-KO HAP1 cells. **(A)** Differentially expressed genes (DEGs) identified in the RNA-seq analyses of MNT-KO (left) or CCDC6-KO HAP1 (right) cells compared to parental (PAR) cells. Genes were clasified as upregulated log2FC ≥ 1 (blue) or downregulated log2FC ≤ –1 (red) with adj.pvalue ≤ 0.05. **(B)** Validation of RNA-seq results in CCDC6-KO HAP1 cells. Log2FC values obtained from RNA-seq analyses (light grey) were compared with RT-qPCR measurements normalized to *RPS14* (dark grey). Error bars represent ±SD from three independent experiments. **(C)** Hallmark pathway enrichment analysis was performed using GSEA on ranked gene expression profiles from MNT-KO and CCDC6-KO cells compared with parental (PAR) controls. The top positively and negatively enriched Hallmark gene sets (FDR < 0.25) are shown together with their normalized enrichment score (NES) and adjusted p-values. **(D)** Enrichment plots for the HALLMARK EPITHELIAL MESENCHYMAL TRANSITION gene set in MNT-KO and CCDC6-KO HAP1 cells relative to parental (PAR) controls. Analyses were performed using the MSigDB Hallmark gene set collection (h.all.v2024.1.Hs).

**SUPPLEMENTARY FIGURE 3.**
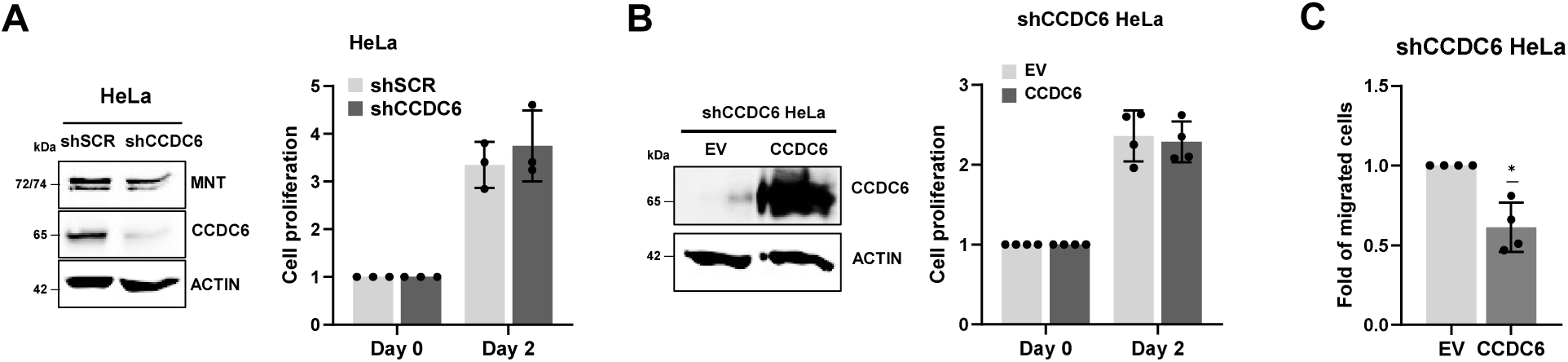
CCDC6 regulates cell migration, but not proliferation, in HeLa cells. **(A)** Validation of CCDC6 silencing in HeLa cells by western blot. CCDC6 and MNT protein levels are shown in HeLa cells constitutively expressing a shRNA against CCDC6 (shCCDC6) or a scrambled control shRNA (shSCR). β-ACTIN levels were used as loading control (left). Cell proliferation was assessed by crystal violet staining (right). Data are presented as mean ± SD from n = 3 independent experiments. Statistical significance was determined using a two-tailed t-test comparing shCCDC6 and shSCR cells at 48 h; ns p > 0.05. **(B)** Validation of CCDC6 overexpression in HeLa shCCDC6 cells by western blot. CCDC6 protein levels are shown in HeLa shCCDC6 cells transduced with CCDC6 or the corresponding control vector. β-ACTIN levels were used as loading control (left). Cell proliferation was assessed by crystal violet staining (right). Data are presented as mean ± SD from n = 4 independent experiments. Statistical significance was determined using a two-tailed t-test comparing EV and CCDC6 cells at 48 h; ns p > 0.05. **(C)** Quantification of transwell migration assays performed with shCCDC6 HeLa cells overexpressing CCDC6 relative to the corresponding empty vector. Cells were allowed to migrate for 20 h under low-serum conditions. Data are presented as mean ± SD from n = 4 independent experiments. Statistical significance relative to the control (set to 1) was determined using a two-tailed one-sample t-test; * p < 0.05.

**SUPPLEMENTARY FIGURE 4.**
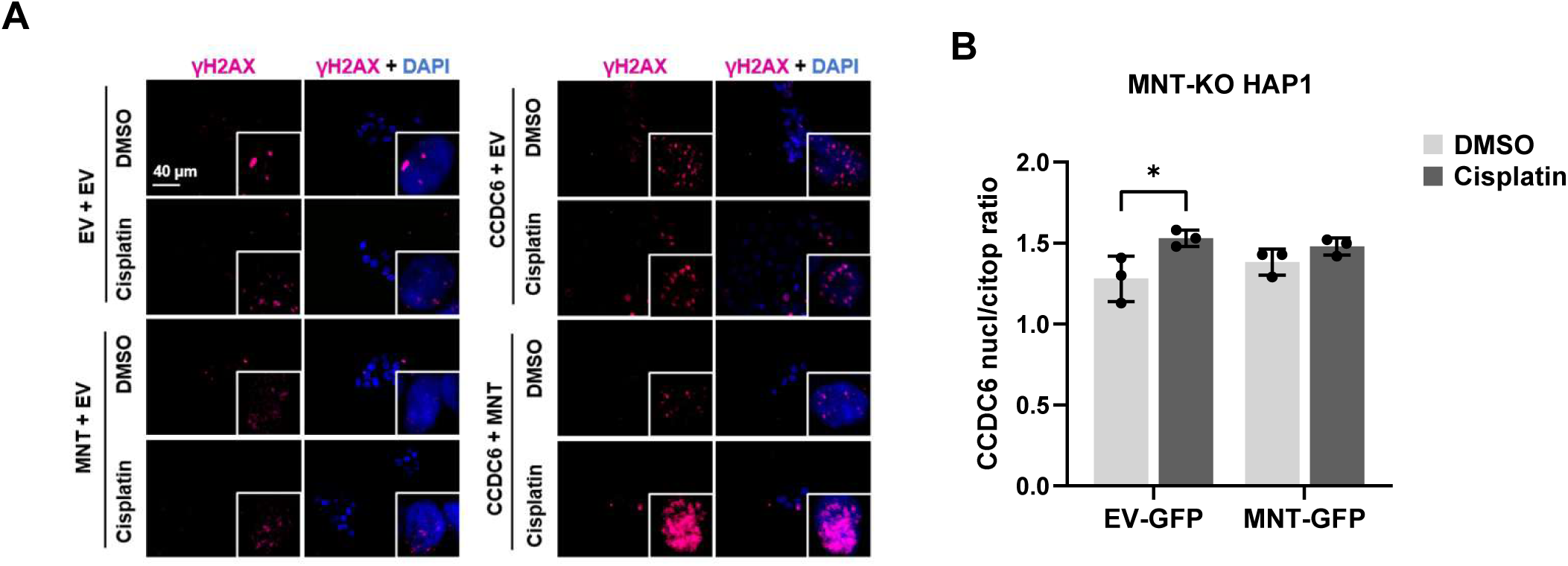
MNT and CCDC6 cooperate in the DNA damage response by modulating γH2AX signalling. **(A)** Representative immunofluorescence images corresponding to the experiments shown in Figures 8E and 8F. γH2AX foci are shown in magenta and nuclei were counterstained with DAPI (blue). MNT-KO HAP1 cells stably expressing CCDC6-Cherry or the corresponding empty vector (EV) and transiently transduced with MNT-GFP or the corresponding empty vector (EV), as indicated, were left untreated or treated with 10 μM cisplatin for 24 h. Enlarged views of representative fields are shown to facilitate visualization of γH2AX foci. **(B)** Quantification of CCDC6 subcellular localization performed by immunofluorescence. MNT-KO HAP1 cells overexpressing CCDC6 with either MNT rescued or expressing a control vector (EV) treated with 10 μM of cisplatin or vehicle (DMSO). Data are presented as mean of the nuclear/cytoplasmic signal ratio of CCDC6 ± SD from n = 3 independent experiments. Statistical significance was determined by two-way ANOVA followed by Fisher’s LSD multiple comparisons test; * p < 0.05.

**Supplementary Table 2.** Cell lines used. (10F: 10% fetal bovine serum). Vectors are described in the Supplementary Table 3

| Cell line | Background | Culture medium | Source |
|---|---|---|---|
| HEK-293T | Human embryonic kidney cell line expressing SV40 T antigen | DMEM-10F | ATCC |
| HeLa | Human cervical cancer | DMEM-10F | ATCC |
| pLKO HeLa | HeLa derivative cell line transduced with pLKO.1 empty vector | DMEM-10F + puromycin | (Merolla et al., 2012) |
| shCCDC6 HeLa | HeLa derivative cell line (constitutively silenced CCDC6) | DMEM-10F + puromycin | (Merolla et al., 2012) |
| shCCDC6 + EV HeLa | shCCDC6 HeLa derivative cell line (control cherry empty vector) | DMEM-10F + G418 + puromycin | This work |
| shCCDC6 + CCDC6 HeLa | shCCDC6 HeLa derivative cell line (constitutive CCDC6-IRES-Cherry vector) | DMEM-10F + G418 + puromycin | This work |
| HAP1 | Human chronic myeloid leukemia | IMDM-10F | Horizon Discovery |
| MNT-KO HAP1 | HAP1 derivative cell line with MNT knockout generated by CRISPR-Cas9 | IMDM-10F | Horizon Discovery |
| CCDC6-KO HAP1 | HAP1 derivative cell line with CCDC6 knockout generated by CRISPR-Cas9 | IMDM-10F | Horizon Discovery |
| MNT-KO + EV-Cherry HAP1 | MNT-KO HAP1 derivative cell line (Control Cherry empty vector) | IMDM-10F + G418 | This work |
| MNT-KO + CCDC6 HAP1 | MNT-KO HAP1 derivative cell line (constitutive CCDC6-IRES-Cherry vector) | IMDM-10F + G418 | This work |
| MNT-KO + EV-Cherry + EV-GFP HAP1 | MNT-KO+EV-Cherry HAP1 transduced with the GFP empty vector | IMDM-10F + G418 + puromycin | This work |
| MNT-KO + EV-Cherry +MNT-GFP HAP1 | MNT-KO+EV-Cherry HAP1 transduced with the MNT-GFP vector | IMDM-10F + G418 | This work |
| MNT-KO + CCDC6 + EV-GFP HAP1 | MNT-KO+CCDC6 HAP1 transduced with the GFP empty vector | IMDM-10F + G418 + puromycin | This work |
| MNT-KO + CCDC6 + MNT-GFP HAP1 | MNT-KO+CCDC6 HAP1 transduced with the MNT-GFP vector | IMDM-10F + G418 + puromycin | This work |
| MCF7 | Human breast cancer cell line | DMEM-10F | ATCC |
| MNT-KO MCF7 | MCF7 derivative cell line with MNT knockout generated by CRISPR-Cas9 | DMEM-10F | This work |
| CCDC6-KO MCF7 | MCF7 derivative cell line with CCDC6 knockout generated by CRISPR-Cas9 | DMEM-10F | This work |

**Supplementary Table 3.** Expression plasmids and lentiviral constructs.

| Name | Construct | Antibiotic resistance | Source |
|---|---|---|---|
| pcDNA 3.1 Zeo | Empty vector | Zeocin | (Liano-Pons et al., 2021) |
| pcDNA 3.1 WT Mnt – HA | Mouse Mnt with HA-tag (C-t) | Zeocin | (Liano-Pons et al., 2021) |
| pcDNA 3.1 ΔNt1 Mnt–HA | Mouse Mnt with HA-tag (C-t) and a deletion of the N-t domain (1-301 aa) | Zeocin | (Liano-Pons et al., 2021) |
| pcDNA 3.1 ΔNt2 Mnt–HA | Mouse Mnt with HA-tag (C-t) and a deletion of the N-t domain (1-273 aa) | Zeocin | (Liano-Pons et al., 2021) |
| pcDNA 3.1 ΔCt1 Mnt–HA | Mouse Mnt with HA-tag (C-t) and a deletion of the C-t and the SID domain (315-591 aa) | Zeocin | (Liano-Pons et al., 2021) |
| pcDNA 3.1 ΔCt2 Mnt–HA | Mouse Mnt with HA-tag (C-t) and a deletion of the C-t (202-591 aa) and the SID domain (1-19 aa) | Zeocin | (Liano-Pons et al., 2021) |
| pcDNA4 WT CCDC6-myc | Human CCDC6 with myc-tag (C-t) | Zeocin | (Celetti et al., 2004) |
| pcDNA4 CCDC6 1-101-myc | 1-101 amino acids of human CCDC6 fragment with myc-tag (C-t) | Zeocin | (Celetti et al., 2004) |
| pcDNA4 CCDC6 1-223-myc | 1-223 amino acids of human CCDC6 fragment with myc-tag (C-t) | Zeocin | (Celetti et al., 2004) |
| Lv103 GFP | Lv103 GFP lentiviral vector (control). | Puromycin | Genecopoeia |
| Lv103 MNT-GFP | Lv103 human MNT with GFP-tag (N-t) lentiviral vector | Puromycin | Genecopoeia |
| pLV Cherry-IRES-neo | pLV-EF1A mCherry lentiviral vector (control) | G418 | Vector Builder |
| pLV CCDC6-IRES-mCherry-IRES-neo | pLV-EF1A-Human CCDC6-IRES Cherry lentiviral vector | G418 | Vector Builder |
| LentiCRISPRv2GFP-MNT | LentiCRISPRv2GFP containing gRNA for MNT knockout | Puromycin | Addgene |
| LentiCRISPRv2GFP-CCDC6 | LentiCRISPRv2GFP containing gRNA for CCDC6 knockout | Puromycin | Addgene |
| pMDL | Packaging plasmid containing the HIV gag-pol gene | None | Addgene |
| pREV | Packaging plasmid containing the HIV rev gene | None | Addgene |
| pCMV-VSV-G | VSV-G gene encoding enveloped lentiviral protein | None | Addgene |
| psPAX2 | GAG and POL genes encoding packaging lentiviral proteins | None | Addgene |

**References**
Celetti, A., Cerrato, A., Merolla, F., Vitagliano, D., Vecchio, G., and Grieco, M. (2004). H4(D10S170), a gene frequently rearranged with RET in papillary thyroid carcinomas: functional characterization. Oncogene *23*, 109-121.
Liano-Pons, J., Lafita-Navarro, M.C., Garcia-Gaipo, L., Colomer, C., Rodriguez, J., von Kriegsheim, A., Hurlin, P.J., Ourique, F., Delgado, M.D., Bigas, A.*, et al.* (2021). A novel role of MNT as a negative regulator of REL and the NF-kappaB pathway. Oncogenesis *10*, 5.
Merolla, F., Luise, C., Muller, M.T., Pacelli, R., Fusco, A., and Celetti, A. (2012). Loss of CCDC6, the first identified RET partner gene, affects pH2AX S139 levels and accelerates mitotic entry upon DNA damage. PloS one *7*, e36177.

**Supplementary Table 4.**
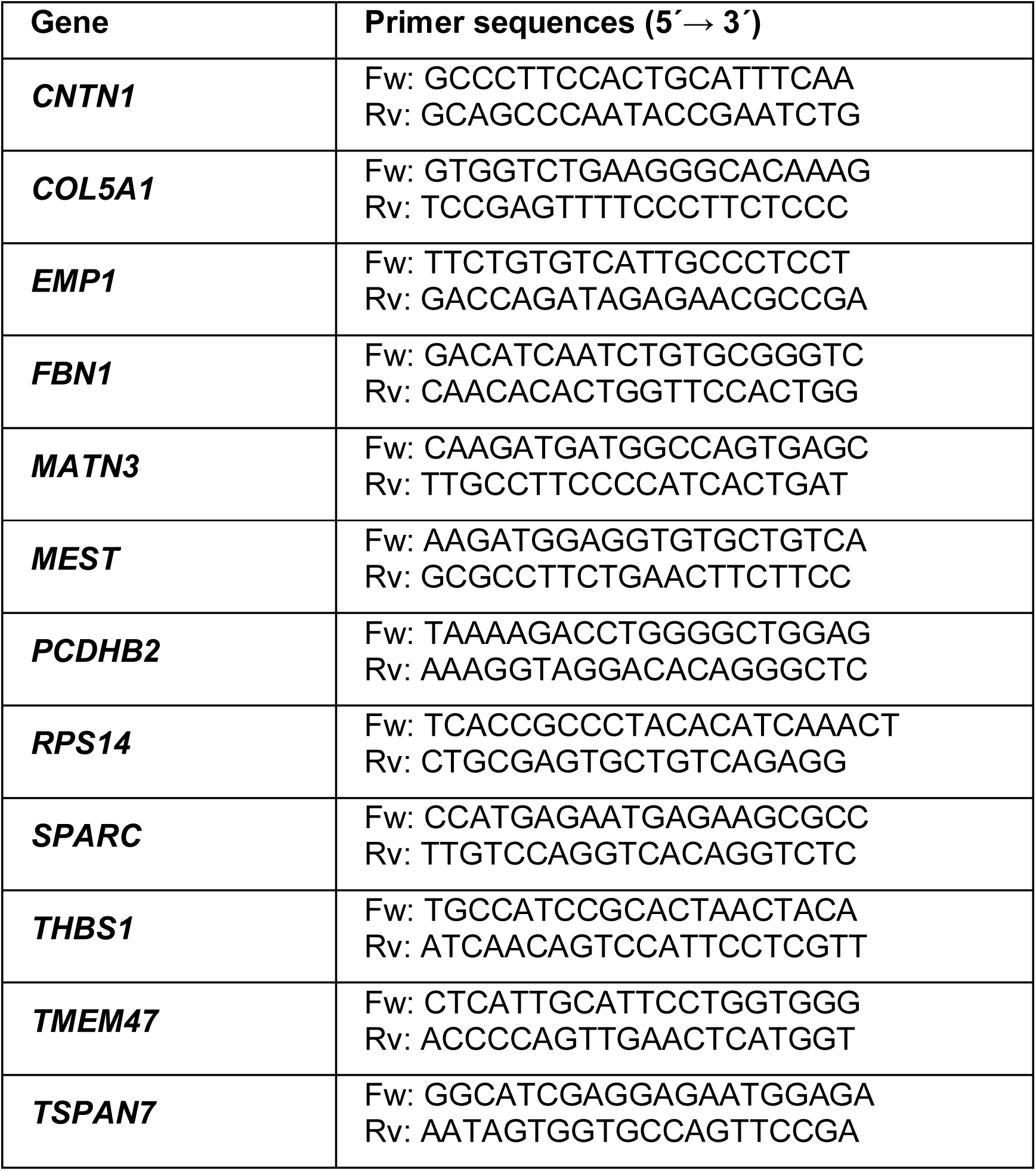
Primers used for RT-qPCR. Fw: forward primer; Rv: reverse primer

**Supplementary Table 5.** Primary antibodies. IF, immunofluorescence; IP, immunoprecipitation; WB: western blot.

| **Primary antibody** | **Specie and type** | **Reference** | **Technique (dilution)** |
|---|---|---|---|
| β-ACTIN | Mouse monoclonal | Santa Cruz Biotech. (C-4, sc-47778) | WB (1:3,000) |
| β-tubulin | Mouse monoclonal | Santa Cruz Biotech. (D-10, sc-5274) | WB (1:1,000) |
| CCDC6 | Mouse monoclonal | Santa Cruz Biotech. (Q-23, sc-100309) | WB (1:1,000); IF (1:500) |
| CCDC6 | Rabbit polyclonal | Cohesion Biosciences (QA3990) | WB (1:2,000); IF (1:200) |
| GFP | Rabbit polyclonal | Invitrogen (A-11122) | WB (1:1,000); IF (1:200) |
| ERK2 | Mouse monoclonal | Santa Cruz Biotech. (D-2, sc-1647) | WB (1:1,000) |
| pERK2 | Mouse monoclonal | Santa Cruz Biotech. (E-4, sc-7383) | WB (1:1,000) |
| γH2AX | Mouse monoclonal | Millipore (05-636) | IF (1:500-1:1,000) |
| HA | Mouse monoclonal | Genecopoeia (CGAB-HA-0050) | IP |
| MAX | Rabbit polyclonal | Santa Cruz Biotech. (C-17, sc-197) | WB (1:1,000) |
| MNT | Rabbit polyclonal | Novus (NBP2-04052) | IP |
| MNT | Rabbit polyclonal | Novus (NBP2-04053) | WB (1:1,000) |
| MYC | Mouse monoclonal | Santa Cruz Biotech. (9E10, sc-40) | IP, WB (1:1000) |
| THBS1 | Mouse monoclonal | Santa Cruz Biotech (C-8, sc-393504) | WB (1:1000) |

**Supplementary Table 6.** Secondary antibodies. IF, immunofluorescence; WB: western blot.

| **Secondary antibody** | **Specie and type** | **Reference** | **Technique (dilution)** |
|---|---|---|---|
| Anti-Mouse IRDye®680 | Donkey polyclonal | LI-COR (926-68072) | WB (1:10,000) |
| Anti-Mouse IRDye®800 | Donkey polyclonal | LI-COR (926-32212) | WB (1:10,000) |
| Anti-Rabbit IRDye®680 | Donkey polyclonal | LI-COR (926-68073) | WB (1:10,000) |
| Anti-Rabbit IRDye®800 | Donkey polyclonal | LI-COR (926-32213) | WB (1:10,000) |
| AlexaFluor 647-conjugated anti-mouse | Donkey polyclonal | Invitrogen (A31571) | IF (1:200-1:500) |

